# Coupled dynamics of charged macromolecules and counterions mediated by binding enzymes

**DOI:** 10.1101/2021.06.21.449292

**Authors:** Tapas Singha, Siao-Fong Li, Murugappan Muthukumar

## Abstract

We investigate the role of active coupling on the transport properties of the macromolecules. The active coupling comes due to bound enzymes with a segment of the macromolecule wherein the enzyme exerts an electrostatic force on the segment of the macromolecule, and eventually, it gets unbound due to the thermal fluctuations. This binding and unbinding process generates active fluctuations in the dynamics of the macromolecule. Starting with segment dynamics and correlations for three dynamical models with active coupling, we obtain the cooperative diffusivity for the realistic charged macromolecules with hydrodynamics. First, we construct the three models by incorporating the features of a real polymer systematically, starting from simple Rouse dynamics with active coupling. We further include segment-segment interactions and in addition, hydrodynamic interactions with active coupling. Our obtained scaling form for segment-segment correlations for the models in terms of the size exponent of the polymer indicating that hydrodynamic and segment-segment interactions along with the active coupling lead to new scaling regimes. We finally study the dynamics of a homogeneously charged flexible polymer in an infinitely dilute solution where enzymes and counterions affect the dynamics of the polymers. We analytically investigate how these active fluctuations affect the coupled dynamics of the polymer and counterions. It turns out that these active fluctuations enhance the effective diffusivity of the polymer. The derived closed-form expression for diffusivity is pertinent to accurate interpretation of light scattering data on multi-component systems with binding-unbinding equilibria.

## I. INTRODUCTION

A widespread biological system consisted of macromolecules namely nucleotides, proteins, enzymes, etc., carry electric charges. The dynamics of these charged macromolecules in a solution, are typically governed by several physical and chemical driving forces, and thereby the transport properties of the macromolecules are also affected by these forces. There have been a significant number of biophysical processes such as dynamics of the molecules due to protein binding, the action of the myosin on actin filaments in a cytoskeleton network [15], remodeling of chromatin by ATPase [16] etc., which strongly depend on the transport properties of the macromolecules. Besides, the biological systems, in a synthetic polymeric system, for biomedical application such as drug delivery, to use of microdevice and nanomedicine [6–8], the understanding and controlling the dynamical properties are essential requirements.

In the last two decades, there has been a growing interest in the field of macromolecules (or polymers) which consumes energy from ATP hydrolysis, in which, the macromolecules are either consisted of active monomers [21] or passive polymer embedded in a bath of self-propelled active particles [23, 24]. A large number of theoretical and numerical studies have been carried out starting from the dynamics of a single polymer [1, 2, 26, 28], to a collection of polymers driven by active coupling [3], in which various aspects of polymers such as activity-induced swelling [1, 22, 25], aggregation[20], and patterns [19], of the polymers, have been investigated.

A theoretical study of self-propelled colloidal particles shows that diffusivity due to self-electrophoresis [34] get increased and experiments of catalytic enzymes demonstrate [35, 36] the enhancement of the diffusivity via electrophoresis. In other experiments on transcription of RNA polymerase on a Fluorescently labeled DNA template [30] demonstrate that the diffusion coefficient of DNA gets enhanced significantly. Though the enzyme’s diffusivity has been studied in the presence of substrate, the studies of the effect of enzyme’s on macromolecule have yet to be explored. In other theoretical study, a neutral flexible polymer in an environment of active enzymes (energized by ATP) is considered where enzymes generate dipolar force along the polymer backbone upon binding [27]. However, as per our knowledge, the study of polymer, hardly considers the charged species explicitly and its electrostatic interactions as a source of activity, though most of the cellular components are charged, and induce active coupling upon binding *without* ATP hydrolysis. This type of active coupling is basically different from the usual active particle which typically consumes energy from ATP hydrolysis and dissipates energy by exerting mechanical force [3].

On the other hand, there has been a vast area in the field of charged macromolecules wherein dynamical properties for a charged polymeric solution have extensively been studied in a number of theoretical, numerical, and experimental work [4, 5]. Though the solution as a whole is electrically neutral, the dynamics of a charged polymer in a solution are strongly coupled by the electrostatic interactions of the dissociated counterion’s dynamics.

In this view, as on one hand, the ATP hydrolysis mediated active dynamics of macromolecules have hugely been studied in active matter, on the other hand, electrostatic driven macromolecular dynamics have been studied extensively, still, electrostatic interactions may induce active coupling without ATP hydrolysis in this charged systems that have not been realized so far in both the fields. This work may bridge between these two growing fields.

In particular, in order to understand the effect of active coupling in the dynamics of the polymer in a systematic way, we study the three dynamics for the macromolecules and investigate static segment-to-segment correlation and mean-squared displacement of the segments. We start with simple Rouse dynamics with active coupling where hydrodynamic interactions and segment-segment interactions are neglected. Thereafter, we incorporate the segment-segment interactions but not hydrodynamic interactions which is defined as Generalized Rouse dynamics (GR). In the third system, we also include hydrodynamic interactions along with the segment-segment interactions, Generalized Zimm dynamics (GZ) and study the effect of active polymer on static and dynamic correlations. In the segment-segment interactions, we consider the excluded volume and the electrostatic interactions between all possible pairs of segments.

Finally, we aim to understand the transport properties of a charged macromolecule in an infinitely dilute solution with active coupling where activity arises due to electrostatic interactions between segments and protein-like enzymes. To do so, in this setting, first of all, there has been frictional force between charged segments and solvent, mobility of the polymer in the presence of counterions are to be taken care of. Following refs. [4, 37] we incorporate the translational friction coefficient, mobility of the polymer. Further, we follow the Fokker-Planck formalism for a Brownian particle with colored noise[9] and employ it to obtain the concentration equation of a polymer in the presence of bound enzymes which eventually gets detached and gives rise to a colored noise, and finally we couple the time evolution of the counterion’s concentration and derive the cooperative diffusivity of the polymer in the large scale limit which can be measured in the dynamic light scattering experiment.

We find that the coupled dynamics of the charged polymer and counterions gets significantly affected when the enzymes get actively coupled with the dynamics of the polymer. This active coupling enhances the diffusivity of the charged polymer which is not obvious, because, an enzyme, increases the mass of the composite upon binding than an unbound individual, (and the instantaneous averaged force on the polymer is zero), the diffusivity of the polymer gets enhanced due to this binding process. This happens because the enzyme’s bound process increases the fluctuations in the velocity of the polymer and thereby increases the diffusion constant of the polymer. The systematic understanding of enhancement of the diffusivity of macromolecules may have immense applicability in terms of controlling the transport properties in the biophysical systems.

Considering the counterion’s relaxation is very fast compared to that of the macromolecule’s concentration fluctuations, we obtain a closed-form expression for the cooperative diffusivity which explicitly shows how the strength of the electrostatic force, temperature, and screening length controls the diffusivity. This may be very useful for future experimental measurements.

We first start with a segmental dynamics for three different models which are considered incorporating features of charged polymeric model systematically. Then we study the cooperative diffusivity of polymer in a dilute solution. Here we organize our work in the following way. In Section-II, we define the three models and describe the basics features of the models. In Section-III, we present the time independent segment-segment correlation for all three models. In Section-IV, we present MSD for the models. Next we focus on the cooperative diffusivity of a macromolecule in Section-V. The section-VI, presents the coupling of the concentration fluctuations of the counterions and polymer. Finally, we have obtained cooperative diffusivity of the charged polymer for this active coupling in dilute solutions. In Section-VII, we summarize our results and conclude our work.

## II. DYNAMICAL MODELS OF MACROMOLUCULES WITH ACTIVE COUPLING

We first start with simple Rouse dynamics with active coupling wherein we consider a very long polymer where *s* is continuous variable. The dynamical equation for a segment can be written as

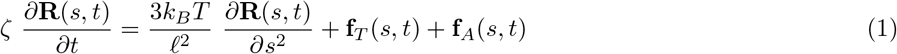

where *ζ* is friction coefficient of a segment, and *ℓ* is Kuhn length of the polymer. **R**(*s, t*) is position vector of sth segment, and note that *s* is the dimensionless index of a segment along the contour. The first term on the right hand side due to connectivity, and the second term *f_T_*(*s, t*) is the thermal noise which is Gaussian with 〈*f_T_*(*s, t*)〉 = 0 and auto-correlation of one component of the noise **f**_*T*_(*s, t*) is the following

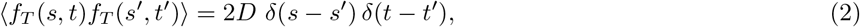

and the third term represents the bound and unbound process of the enzymes described in detail below.

### A. Autocorrelation of active force

The term **f**_*A*_(*s, t*) in Eq.(1) is the force exerted on a segment due to actively binding and unbinding of an enzyme to a polymer (shown in Fig. 1). The ensemble average of the force of any component of the term can be considered to be zero. When a single enzyme and a segment of a polymer participate in the binding and unbinding process, each bound event is followed by an unbound event, and in an unbound state, the amplitude is zero. Considering the kinetics of the bound and unbound dynamics of the enzyme, which is a telegraphic process, an auto-correlation [18] can be obtained as

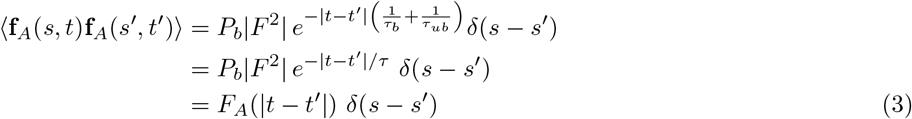

where 〈…〉 indicates ensemble average where *τ_b_* and *τ_ub_* are the average bound and unbound time. In this work, we consider the enzymes with the charged polymeric solution are in thermal equilibrium. Thus at time *t*, the probability of a segment to be bound to an enzyme is

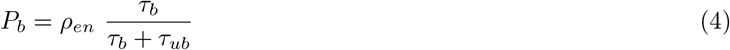

**FIG. 1.**
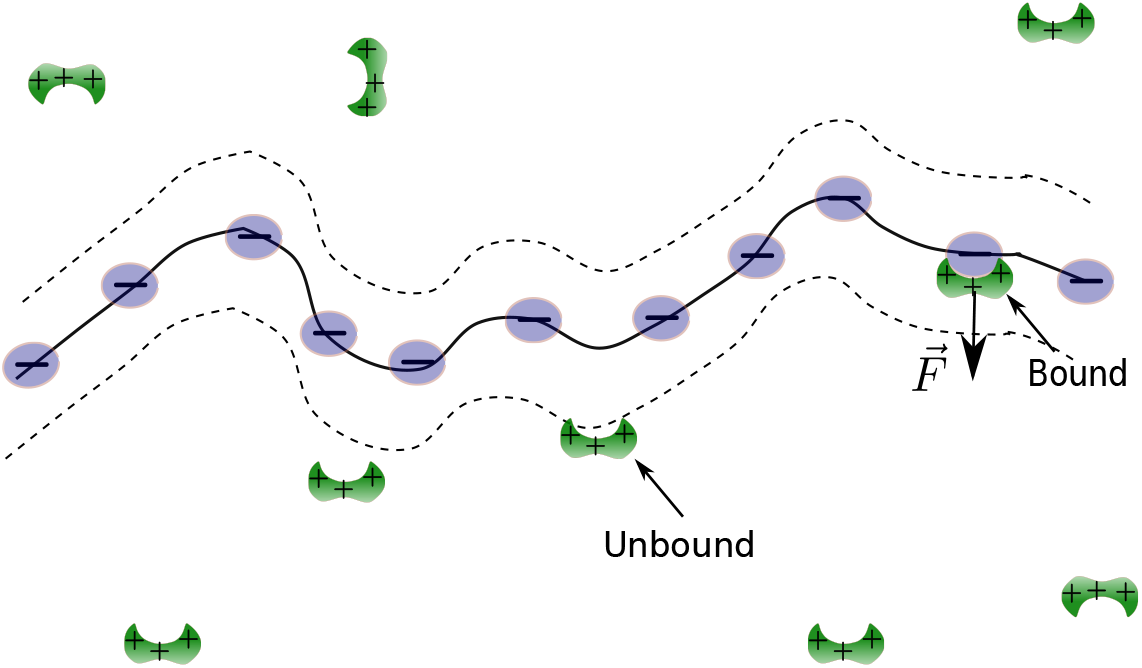
A schematic diagram of a polymer which is negatively charged and enzymes.

Since the enzymes bound due to the electrostatic force, the average bound time can merely be considered as

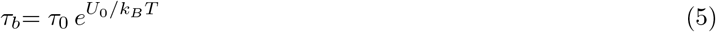

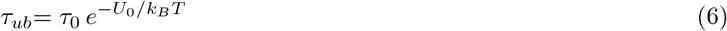

Therefore, probability of binding can also be expressed as

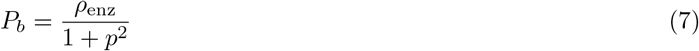

where *p* = *e*^−*U*_0_/*k_B_T*^. The relations are expressed in terms of *U*_0_ indicating potential at *U*(*r*)|_*r*=*xℓ_B_*_ = *U*_0_ where *x* could be fraction or a whole number. For simplicity, we can set it to be *x* = 1

### B. Fourier transformation

We define the convention of the Fourier transformations as

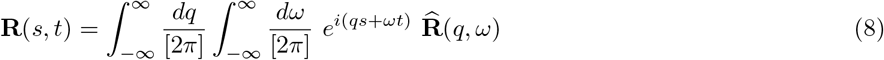

and

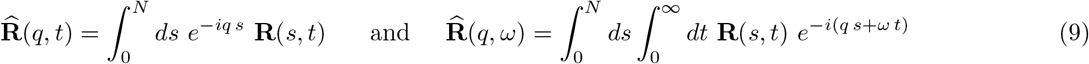

Using the above conventions, autocorrelation of thermal noise (Eq. 2) can be written in Fourier-space as

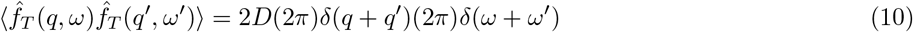

and similarly the auto-correlation of the active-force (Eq. (3)) as

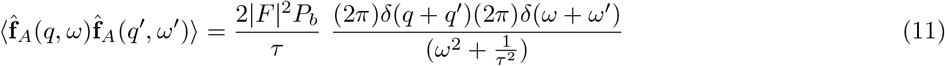

where 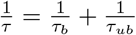.

#### 1. Rouse dynamics with active coupling

Using Eq.9 and 8, the active-Rouse equation can be written in Fourier-space as

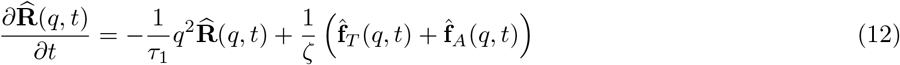

Let us define single-segment’s relaxation timescale as

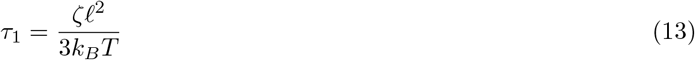

We use the notation *τ*_1_, single segment relaxation timescale of a segment in Rouse chain. In the next subsections, we describe the three models for polymer dynamics wherein qualitatively different features are captured. Apart from the active coupling, the main differences among the three models are the following. For GR model, segment-segment electrostatic and excluded volume interactions are involved along with the connectivity whereas the degree of complexity is greater in the GZ model wherein hydrodynamics interactions are involved in the system.

#### 2. Generalised Rouse dynamics with active coupling

In this model, we consider homogeneously charged macromolecules wherein the segments of the macromolecule interact electrostatically in addition to that there is the excluded-volume interactions. In real space, the equation of motion of a segment can be written as

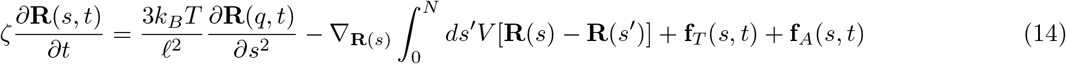

The second term on the right-hand side of the above equation represents the coupling due to the interactions of the sth segment to all oher segments which are highly nonlinear. In the Fourier-space, the mode from third term is coupled to all other modes which is approximately equivalent to the assumption of uniform expansion of the polymer chain with an effective Kuhn length *ℓ*_eff_ [33], and effect of third term modifies the Rouse mode in the second term which is re-expressed in the equation below. In this scenario, the coupling of active force make the dynamics more interesting. The equation of motion of a segment is written in Fourier space as

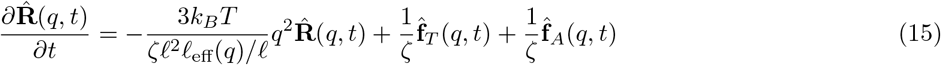

where *ℓ*_eff_(*q*) captures the segmental electrostatic interactions and expressed as *ℓ*_eff_(*q*)/*ℓ* ≃ *a_ℓ_*|*q*|^1−2*ν*^. Thus *a_ℓ_* contains the quantitative information of the interactions and the exponent *ν* contains the conformational information of the polymer. Keeping consistency with the active rouse model, the single-segment’s relaxation timescale can be expressed as

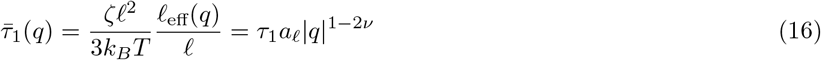

This model captures electrostatic segment-segment interactions (*ℓ*_eff_/*ℓ* ≃ *a_ℓ_*|*q*|^1−2*ν*^) in addition to the connectivity and active coupling as AR. For *ν* = 1/2, the electrostatic interactions are totally screened, and for *a_ℓ_* = 1, where excluded volume interactions is taken to be zero, the GR model essentially becomes the Rouse with active coupling.

#### 3. Generalised active Zimm dynamics

We are ultimately interested in the charged polymeric solution where hydrodynamic interactions becomes very important. In an infinitely dilute solution, the Langevin equation for a charged polymer segment driven by the thermal noise and actively binding and unbinding of enzymes can be expressed as

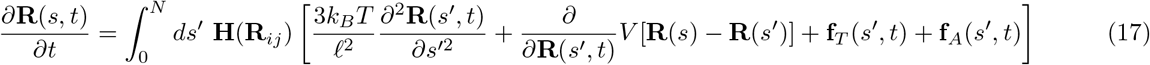

The first term on the right-hand side, represents the conectivity of *s′*th segment, and the interactions between *s*th and *s′*th segments (second term) which affect all other segments of the polymer via hydrodynamics interactions. Similarly, the thermal noise **f**_*T*_(*s, t*) and active coupling **f**_*A*_(*s, t*) affect the whole polymer in the solvent. We first take the Fourier transformation of Eq. (17), and consider pre-averaging approximation for *H*(|**R**(*s*) – **R**(*s*′)|) and uniform expansion approximations [33], yields

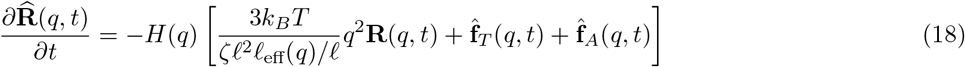

With the hydrodynamic and electrostatic interactions, the single segment timescale can be obtained as

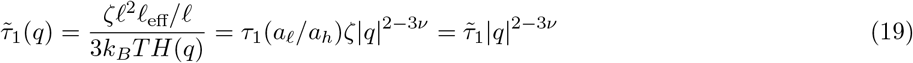

This model captures the hydrodynamics, segment-segment interactions where *H*(*q*) = *a_h_*|*q*|^1−*ν*^. We study the model in detail in the subsequent sections. Since the hydrodynmic interactions becomes unimportant for *ν* = 1, it essentially become GR with active coupling. In our results for different models, these would be an important indicator to check the results.

## III. SEGMENT-TO-SEGMENT CORRELATION FUNCTION

We study the segment-to-segment correlation and MSD of a segment with the active coupling for three polymer models which are qualitatively different. We write segment-to-segment correlation function in Fourier-space as

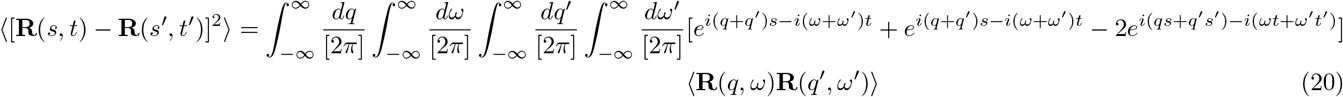

In this subsection, first, we give a few steps for calculating the quantity 〈[**R**(*s, t*) − **R**(*s*′, *t*′)]^2^〉. Expressing the correlation in *q* and *ω* space, we carry out the frequency integrations and keep the form general, so that we can use it for different models. Let us present the segment-segment correlation in Fourier space for active Rouse case as

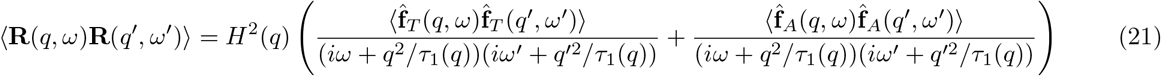

Using the Fourier transformations given by Eqs. 10 and 11, and performing *ω*′ and *q*′ integrations, we obtain

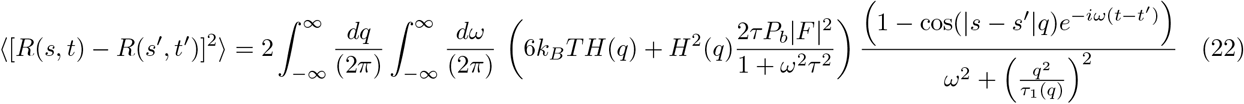

For the sake of presentation, after carrying out frequency integration, we shall also split up the correlation into two parts, namely thermal and active as

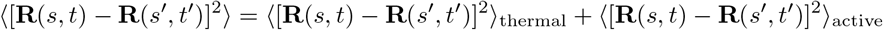

Our obtained thermal and active parts are the following

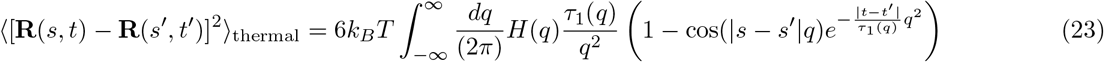

and

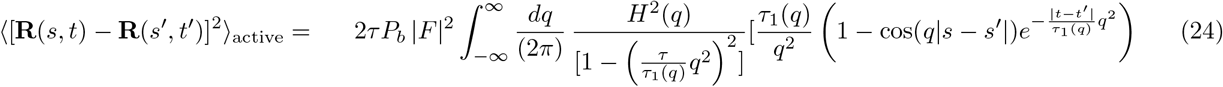

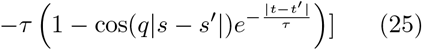

In the above two equations, we present the correlation in terms of general *τ*_1_(*q*) and *H*(*q*) which becomes *τ*_1_ and 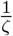 respectively for the Active Rouse model, and *τ*_1_*a_ℓ_*|*q*|^1−2*ν*^ and 1/*ζ* for the generalized-active Rouse models. In the next sections, we present the results of equal-time segment-segment correlation and MSD for the three models and each has thermal and active parts of the correlation. The dynamical structure factor is defined as

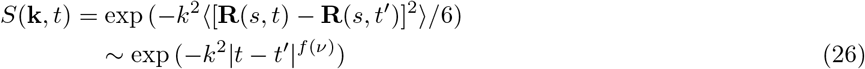

where *k* is the scattering wave vector which can be measured via decay rate 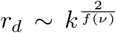 and can be measured in incoherent inelastic neutron scattering.

### A. Time independent segment-segment correlation

We study the time independent segment-to-segment correlation for all the three models. The contribution from the thermal part is compared with the active coupling.

#### 1. Rouse dynamics with active coupling

We investigate the Rouse dynamics with active coupling. It is important to mention that feasible limits of the timescales when *τ*_1_ ≪ *τ* ≪ *τ_R_*. From Eq. 25, for *t* = *t*′, the equal-time thermal part of the correlation can be obtained as

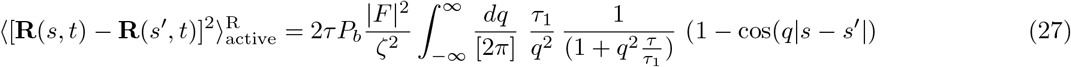

Solving the above integration exactly, we have

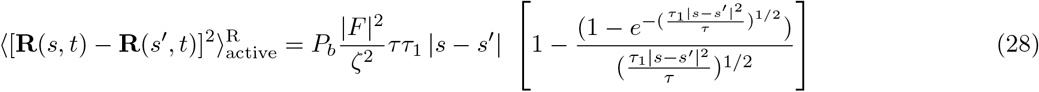

In the asymptotic limits, the equal-time active part of the correlation depends on the limits of the segment’s pair (|*s* – *s*′|) which are following.

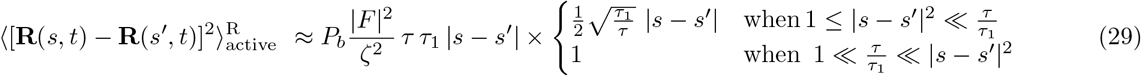

As we see that, the active contribution to the correlation function depends on the ratio of (*τ*_1_/*τ*)|*s* – *s*′| when |*s* – *s*′| is the differences of the segment indices which gets modified by a factor *τ*_1_/*τ* and determines the correlation functions.

The thermal contribution is derived from Eq. (23) as

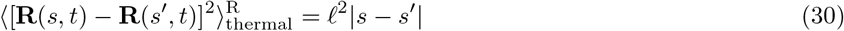

Adding the contribution from thermal and active, we obtain

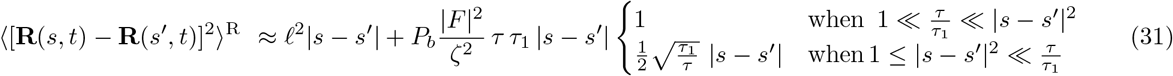

**FIG. 2.**
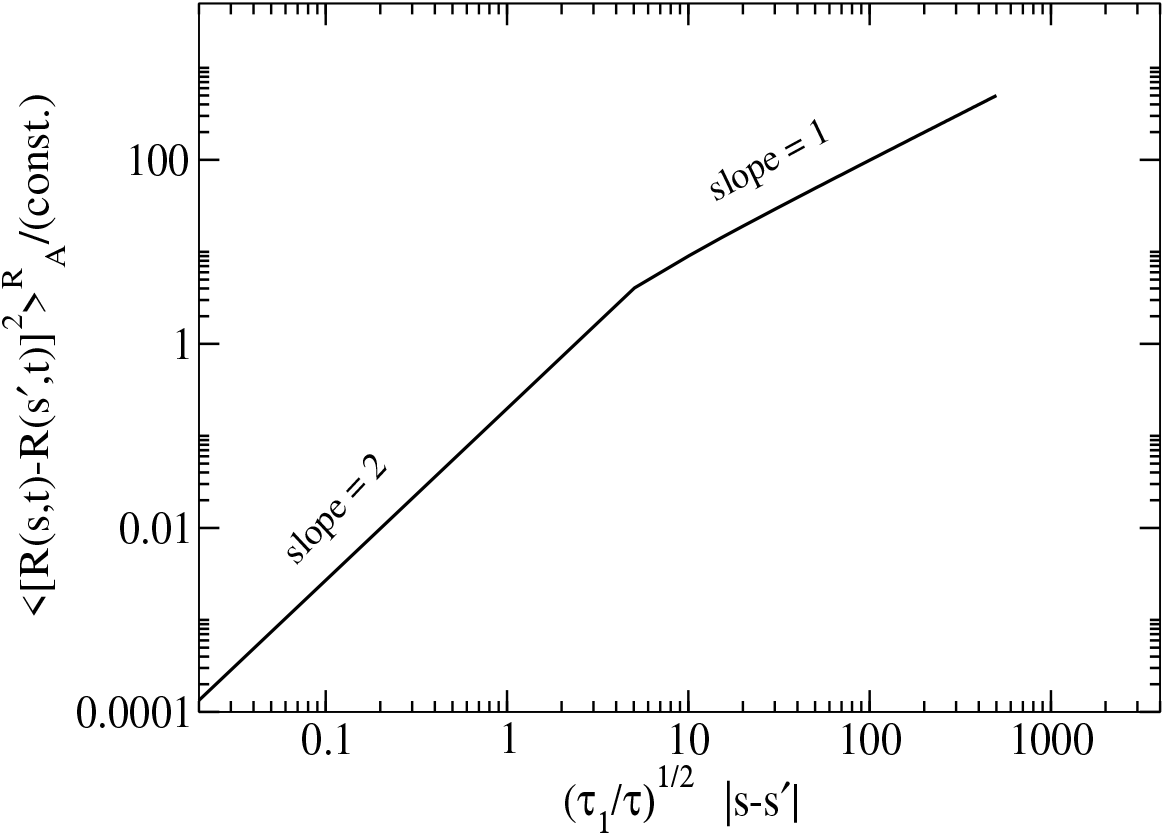
Time independent segment-segment correlation for active-Rouse model. We plot exact results given by the Eq.(28) obtained from the active part which shows that initially it varies |*s* – *s*′|^2^ and in the large scale limit, it behaves as thermal.

We also study equal-time mean-squared end-to-end distance (MSED) which shows MSED gets enhanced by the active coupling for AR and GR models. For GZ model, the scaling law is dominated by active coupling.

#### 2. Mean squared End-to-end distance (MSED)

For moderate *τ*, in the limit *τ*_1_ ≪ *τ* ≪ *τ_R_*, the active part would take large scale limit (|*s* – *s*′| = *N*) from Eq. 29. We obtain the mean-squared end-to-end distance as

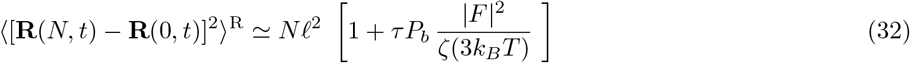

This suggest that equal-time MSED gets enhanced by the active coupling and the increased factor is proportional to the timescale of active coupling and square of the force. Though the magnitude of the force |*F*|, *τ* and *P_b_* depend on the electrostatic interaction which varies with *κ*. Expressing the parameters in terms of *κ*, we observe that the increased factor is dominated by |*F*|^2^ compared to others, and the factor decreases with *κ* because of the exponential relation of |*F*| with *κ*.

#### 3. Generalised Rouse with active coupling

We study the equal-time correlation for GR dynamics with active coupling. We use the general form of the correlation given by Eq. (23). Substituting, *t* = *t*′, and obtain the integral expression for the active contribution of the correlation which reads as

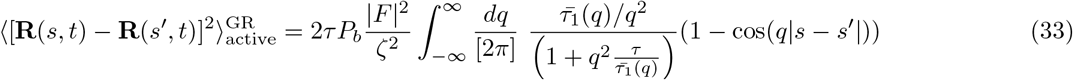

We also substitute the expression of 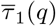 from Eq. 16 in the above equation, we have

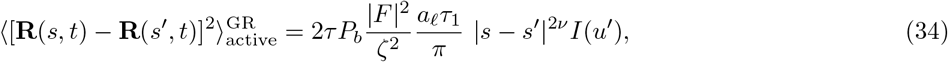

where

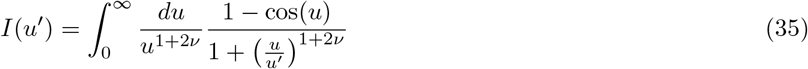

where 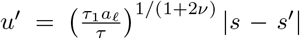 in the above integral. This captures the information of the polymer such as timescale of a single segment relaxation, binding and unbinding of enzymes, information regarding the segment-segment interactions and the lengthscale of the polymer. The above integral fixes the final scaling form of the correlation. We carry out the integral and obtain the results for two limits of *u*′ which are given below.

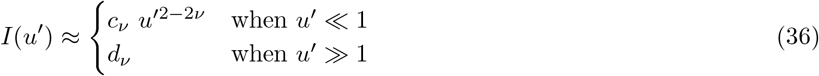

where 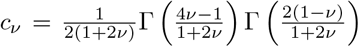 and 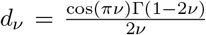. This result can be interpreted in two ways. First approach could be, when the |*s* – *s*′| is fixed but the limit of *τ* is to be set. Second approach, when the *τ* is fixed, for example, *τ*_1_*a_ℓ_* ≪ *τ* ≪ *τ_GR_*. For the second approach, due to the constraints of *τ*, the scaling form is valid only when |*s* – *s*′| satisfies the condition. Substituting Eq. (35) in Eq. 34, we obtain

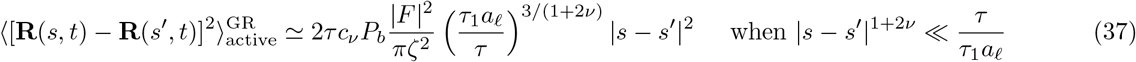

For other limit,

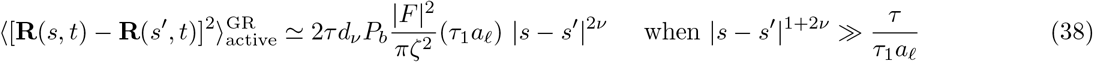

To understand how it varies from one scaling laws to other with *u*′, we also perform numerical integration of Eq. (35) for *u*′ shown in Fig. 3. This numerical evaluation gives insight that how it varies from smaller scale to larger scale. The contribution from the thermal part can be obtained as in Eq. 23, we obtain

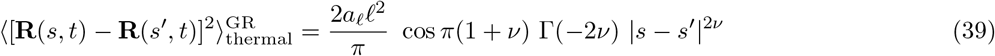

**FIG. 3.**
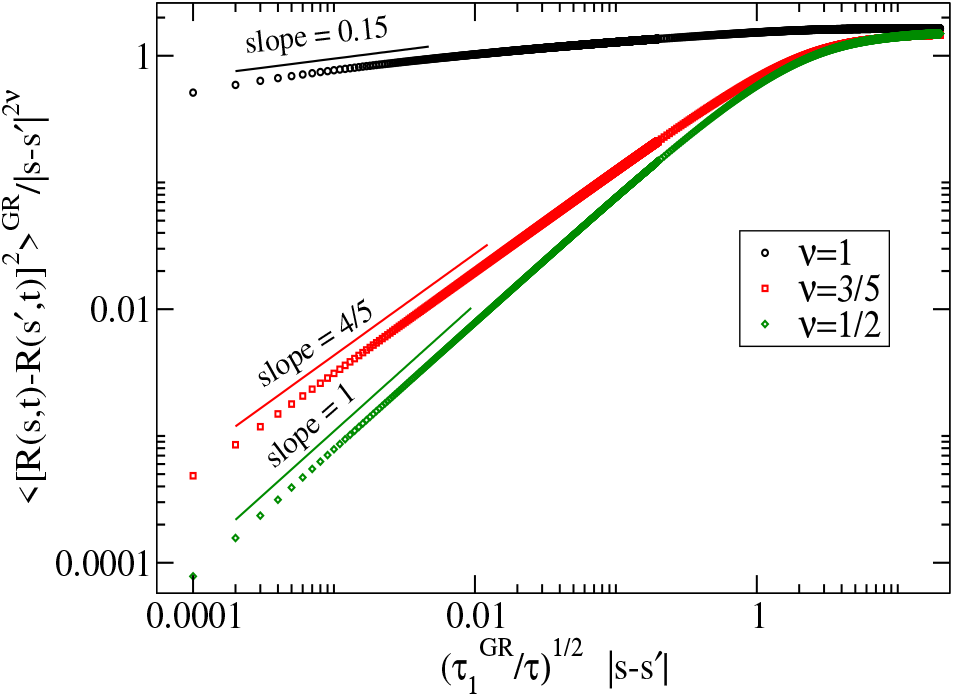
Equal-time segment-to-segment correlation for generalized active-Rouse dynamics where segment-segment electrostatic interactions are considered. We get the data for each value of 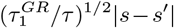 via numerical integration (Eq. (35). This figure shows that the small-scale segment-to-segment distance along the contour swells due to active coupling for a range of *ν* whereas large-scale behavior is thermal as given in Eq. (30). The transition point from rod-like (~ |*s* – *s*′|^2^) to flexible conformation (~|*s* – *s*′|^2*ν*^) is set by the ratio of 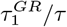. When 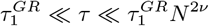, the transition can be observed whereas 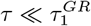, the transition from one scaling to other may not be observed.

By adding the contribution from the thermal and active part for the GR dynamics, we have

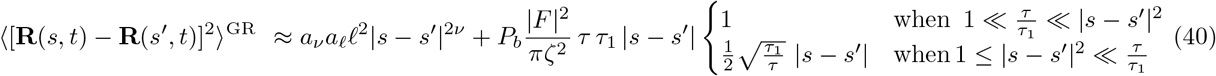

#### 4. Mean squared end-to-end distance

We study equal-time MSED for the GR dynamics where thermal part gets modified by active coupling.

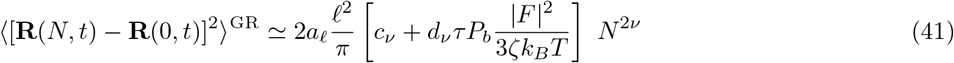

Though the magnitude gets enhanced, the large-scale scaling law of active coupling is the same as its thermal counterpart.

#### 5. Generalised Zimm dynamics with active coupling

Here we study the equal-time segment-segment correlation when the hydrodynamics and electrostatic interactions are considered. We mainly focus on the active part of the correlation and compared with the thermal counterpart. From the given Eq. (25), substituting *t* = *t*′, we obtain

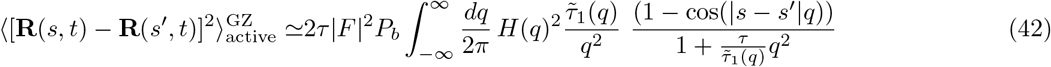

We also substitute the expression for 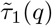 and *H*(*q*) for GZ dynamics in the above equation, and obtain

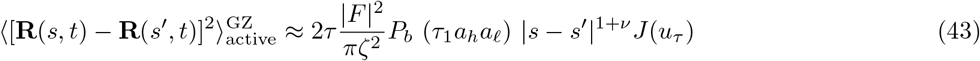

where

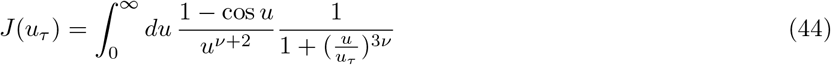

and 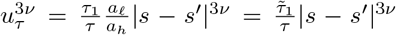. We analytically obtain the scaling for the equal-time correlation in the asymptotic limits.

**FIG. 4.**
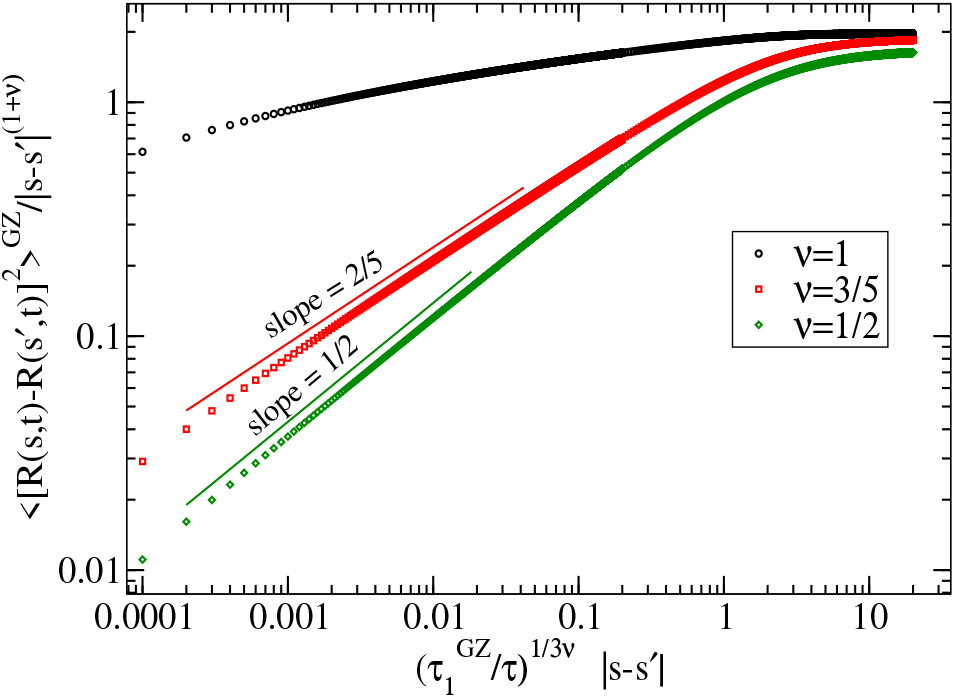
Active contribution to the equal-time segment-to-segment correlation for Generalised-Zimm dynamics with active coupling. Here we plot the results from numerical integration given by Eq. (44). This figure show that the small-scale segment-to-segment distance along the contour behaves rod-like due to active coupling which indicates swelling whereas large-scale behavior is thermal contribution as given in Eq. (46).

The integration over u determines the transition from one limit of *u_τ_* to other limit.

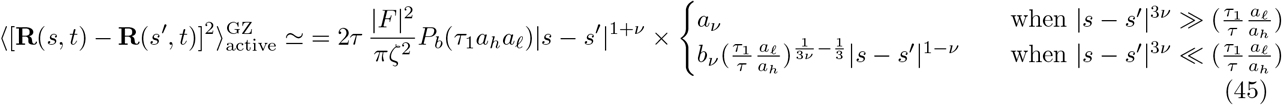

where 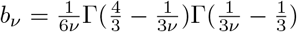 and 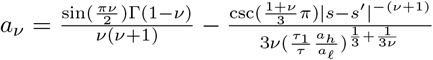.

#### 6. End-to-end distance

We obtain the thermal contribution to the equal-time correlation from Eq. (23) and obtained as

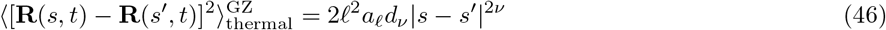

wherein the information of the electrostatic and hydrodynamic interactions are buried in the parameter *a_ℓ_*. For the case of end-to-end distance, with the hydrodynamics interactions, the active force dominates over the thermal noise

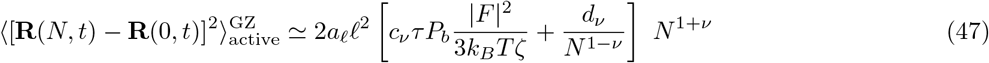

which is evident for large *N*. Therefore, the end-to-end distance varies as *N*^1+*ν*^ in the presence of the hydrodynamics interactions.

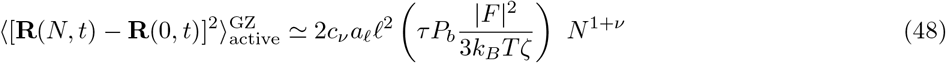

For equal-time correlation, the contribution from the active part show chain swelling (rod-like |*s* – *s*|^2^) when the relative index between two segments is smaller for all the three models whereas for the larger separation the scaling form of the correlation functions converges to its thermal counterpart except for the generalized active Zimm dynamics. In the presence of hydrodynamic interactions, when the relative indices between two segments are large (), the scaling laws of the active part of the correlation do not coincide with the thermal part of the correlation. Therefore, in the large separation limit, the active part (|*s* – *s′*|^1+*ν*^) dominates over the thermal part (|*s* – *s′*|^2*ν*^) which is not the case for the other two models as shown in the Table-1.

**TABLE I.**
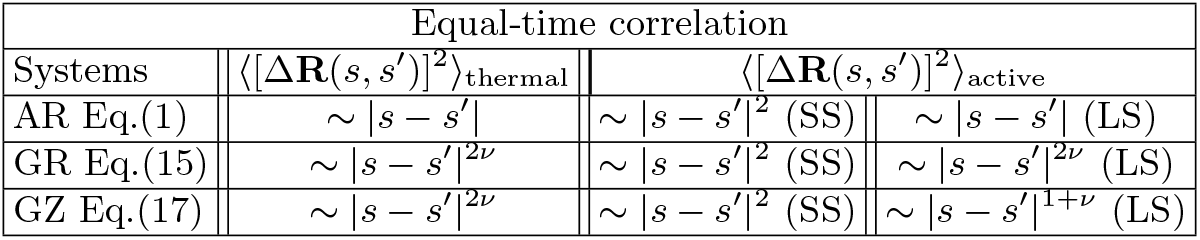
The scaling relations of time independent segment-segment correlation for three dynamics are summarized. In the table, SS, and LS stand for small and large lengthscales, respectively.

For *ν* = 1 the hydrodynamic interactions get totally screened and become scale-independent constant, and the scaling exponent from the active contribution coincides with its thermal counterpart which is expected whereas it deviates for any other values of *ν*. The transitions from smaller segmental separation to larger are shown via numerical integration for each value of scaled separation |*s* – *s′*|. Note that in the asymptotic limits, we obtain the scaling of the active parts of each model with its prefactors, and the results corresponding to the thermal parts are given exactly.

### B. Mean squared displacement

We study the MSD and the scaling forms of a segment for the three models. In addition to the thermal part, the active coupling modifies the correlation significantly in terms of scaling in the short time limit, and the coefficient in the longtime limit. Importantly, in the presence of hydrodynamics interactions the active coupling dominates the scaling in both the limits.

#### 1. Rouse dynamics with active coupling

We study MSD for simple Rouse dynamics with active coupling. To obtain the MSD of a segment, we first substitute 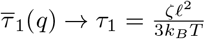 and *H*(*q*) → 1/*ζ*, in Eq. (25) leads to

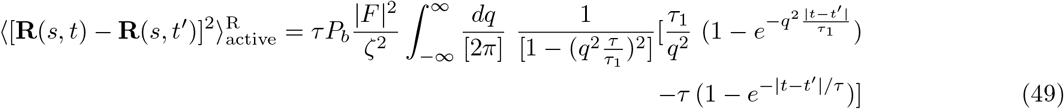

In the above equation, note that there is a divergence in q integration (Eq. 49), we rewrite the integration differently (as shown by Eq. 114 and 115 in the Appendics), so that we can express the *q*th integration independent of |*t* – *t′*|/*τ* After carrying out a few steps (as shown in the Appendics), we obtain

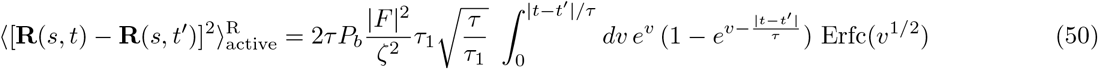

where the integration is performed for full range of *q*. Though the exact solution is obtained for this Active-Rouse model, the expression is complicated to understand the dependencies of |*t* – *t′*|/*τ*. In order to understand the result, we choose to present only the asymptotic limits of the observation time |*t* – *t′*|/*τ* as

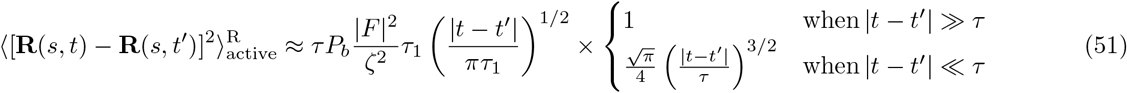

The obtained results in the above equation show Newtonian dynamics in the short time limits (|*t* – *t′*|/*τ* ≪ 1) when the segments do not realize the connectivity whereas in the long time limits the correlation shows ~ |*t* – *t′*|^1/2^ scaling laws similar to the thermal part. We also perform the integration (given by Eq. (50)), numerically, and obtain the asymptotic expressions in the limits of |*t* – *t′*|/*τ*. We compare the results from the numerical integrations with the results obtained from asymptotic limits. The numerical simulation gives us the insight, that how the correlation function depends on |*t* – *t′*|/*τ*. The thermal contribution to the mean-squared displacement of a segment can also be obtained upon substituting *s* = *s′* into Eq. (23).

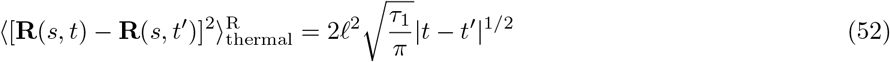

As we mentioned before that the active force brings the coupling between the timescales corresponding to segmental relaxation corresponding to the thermal and active coupling. Depending on the relative magnitude of the timescales, we obtain different results for equal time correlation whereas the thermal part of the correlation is unable to capture these competition of the timescales. Adding the contribution from the thermal and active, we have

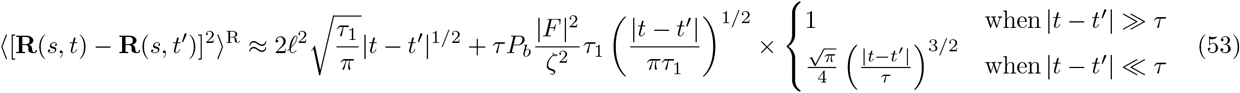

**FIG. 5.**
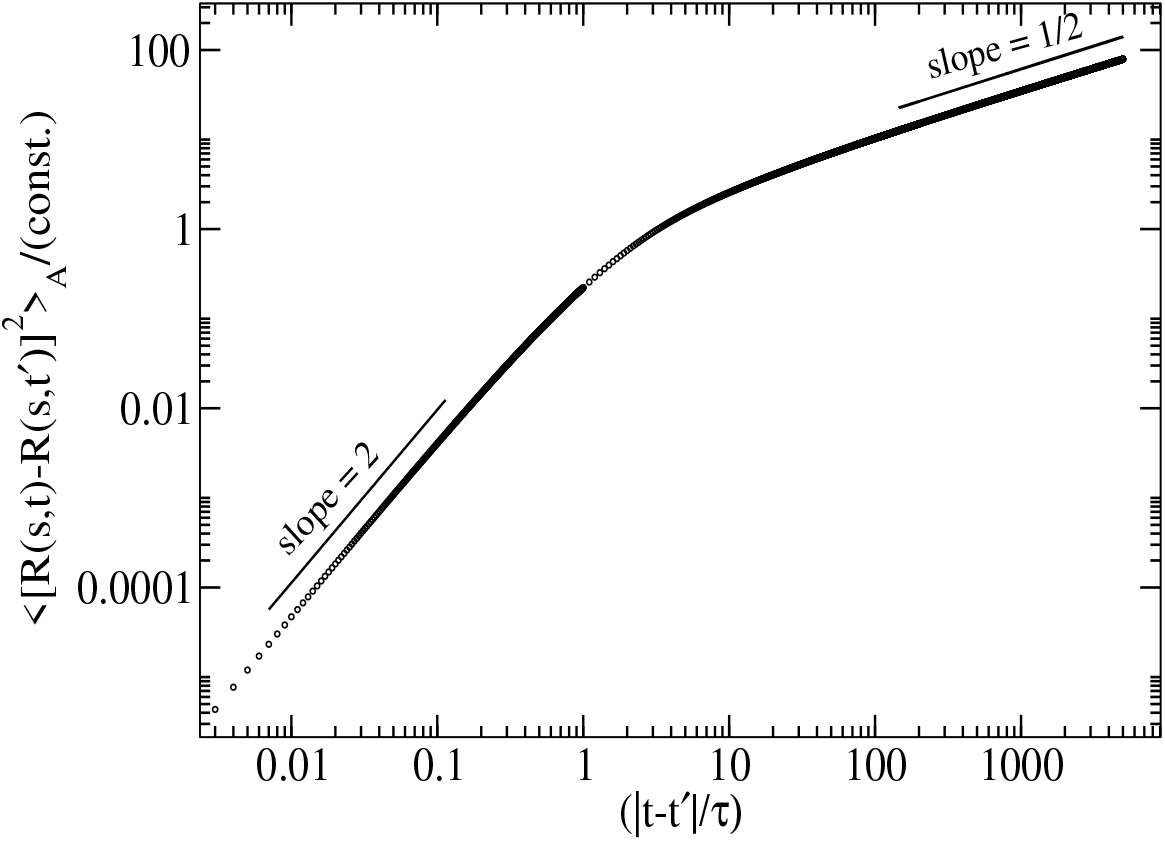
MSD of a segment for Rouse model with active coupling. Here we numerically integrate Eq. 50. This shows how MSD of a segment varies from |*t* – *t′*|^2^ to |*t* – *t′*|^1/2^ for small observation to large observation time limit.

#### 2. Generalised active Rouse

We study the MSD of a segmment in the presence of hydrodynamics and segment-segment interactions. Substituting *s* = *s′* in Eq. 25, the active contribution to the correlation for generalised active Rouse model can be obtained as

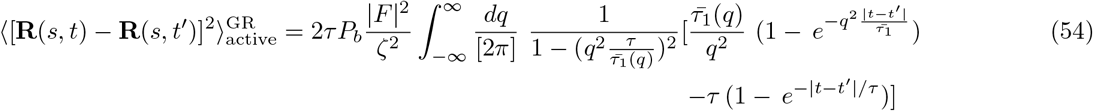

The divergence in *q*th integration is avoided in a non-trivial way (as similar to the calculation as for AR model) and obtain the integral expression as

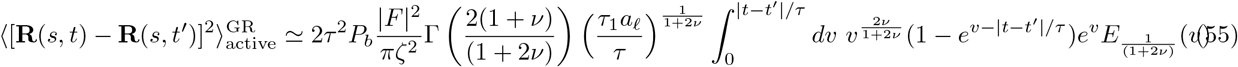

where

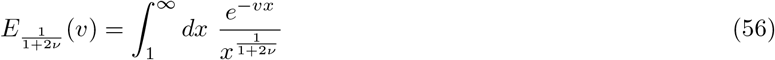

We obtain the analytical results, by considering the asymptotic results of *v* for 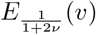. When *v* is small, it suggests that |*t* – *t′*|/*τ* ≪ 1, that leads to

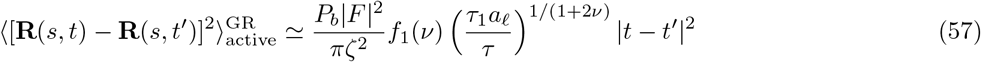

where 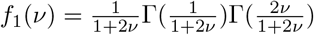. For other limit, for large *v*, i.e., |*t* – *t′*|/*τ* ≫ 1, we obtain

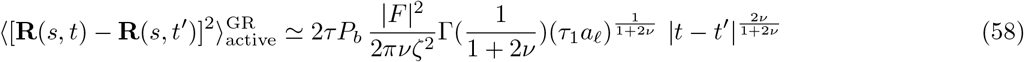

Note that our obtained results from the above integration (Eq. 55) do not depend on the single segment relaxation time 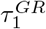. It depends on the relative magnitude of the observation time, |*t* – *t′*| and the the binding and unbinding timescale *τ*. The equation 55 has also been carried out numerically (shown in Fig. 6) which shows that how the correlation function varies from shorter to longer observation time with different scaling exponents. The results from numerical integration agree with the analytical results for each values of *ν* in both the limits. We study the thermal contribution to the MSD, substituting *s* = *s′* in Eq.23 which is obtained as

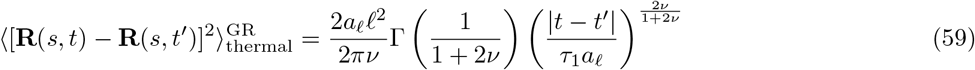

**FIG. 6.**
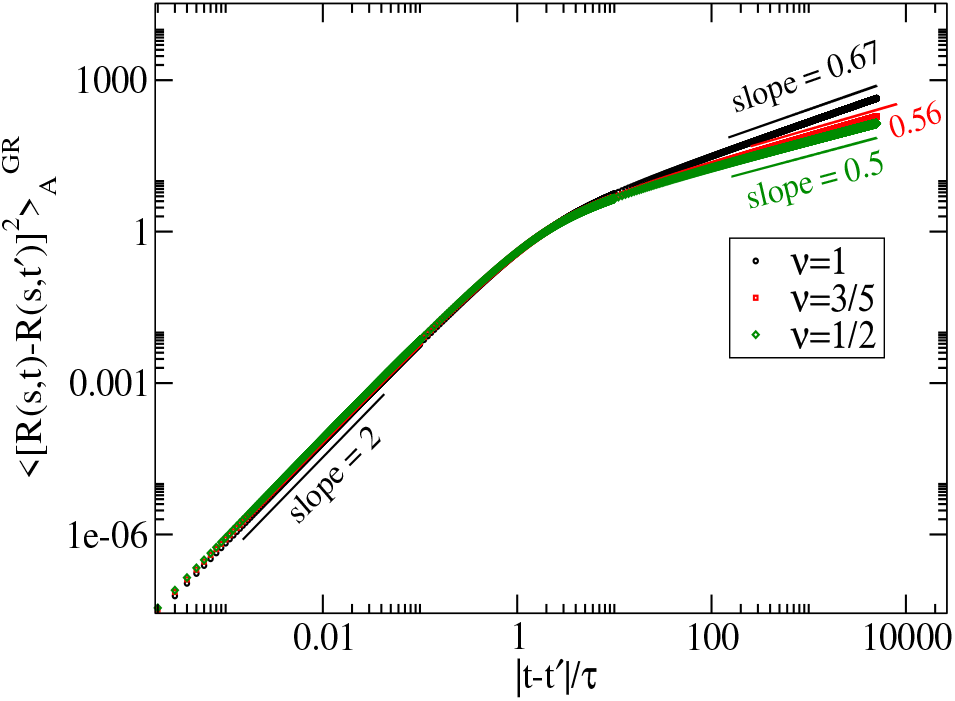
MSD of a segment due to active coupling where segment-segment electrostatic interactions are considered. We numerically evaluate integral expressed in Eq. 55. For short time scale, the correlation shows Newtonian for all range of *ν*. For long observation time, |*t* – *t′*| ≫ *t*, the contribution to the correlation is v independent. This scaling is totally consistent with the asymptotic results from analytics.

Adding the thermal and active part of the above equation, we present the total MSD for GR dynamics. The additional features are captured by GAR than AR is the segment-segment interactions (excluded volume and electrostatic). We consider *ℓ*_eff_/*ℓ* ≃ *a_ℓ_*|*q*|^1−2*ν*^ where *a_ℓ_* contains the information of the excluded volume and electrostatic interactions. The results obtained in Eqs. 57 from the GR will be the same that of AR for *ν* = 1/2 and *a_ℓ_* = 1 which is expected and we also obtained.

The model GR and AR are qualitatively different because GR has the segment-segment interactions, whereas AR does not have segment-segment interactions. The approximated GR dynamics in Fourier space (Eq.15), which we use for the calculation of MSD, can exactly be mapped to AR for *a_ℓ_* = 1 and *ν* = 1/2.

#### 3. Generalised Active Zimm dynamics

We study the time evolution of a segment with hydrodynamic and segment-to-segment interactions. Starting with a Langevin equation, we obtain the corelation given by Eq.25, and substituting *s* = *s′*, we obtain the below equation substituting *s* = *s′* in Eq. 25

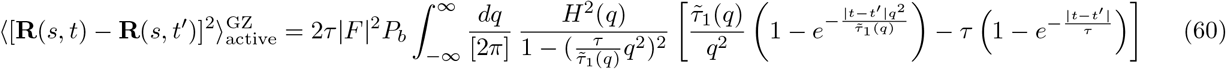

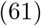

We also substitute the form of *H*(*q*) and 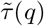 in the above quation, we have

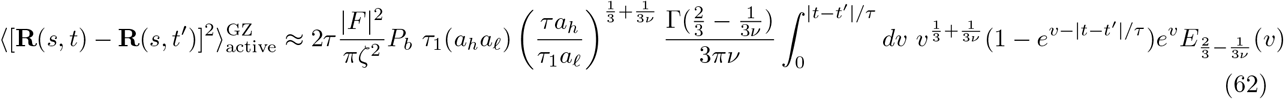

We solve the integration in the asymptotic limits of *v* for the exponential integral *E* which can be obtained as

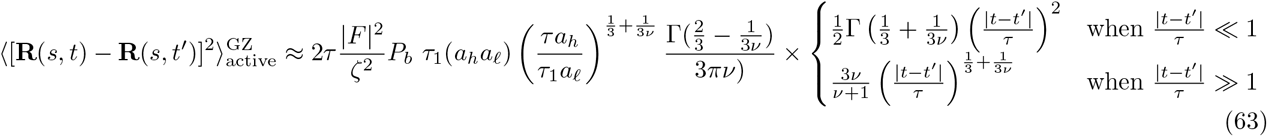

The obtained scaling form is general for any feasible size exponent *ν*. Though in the short time limit, the active part of the correlation varies |*t* – *t′*|^2^, interestingly, the longtime scaling laws does not coincide with the thermal and it is significantly different.

**FIG. 7.**
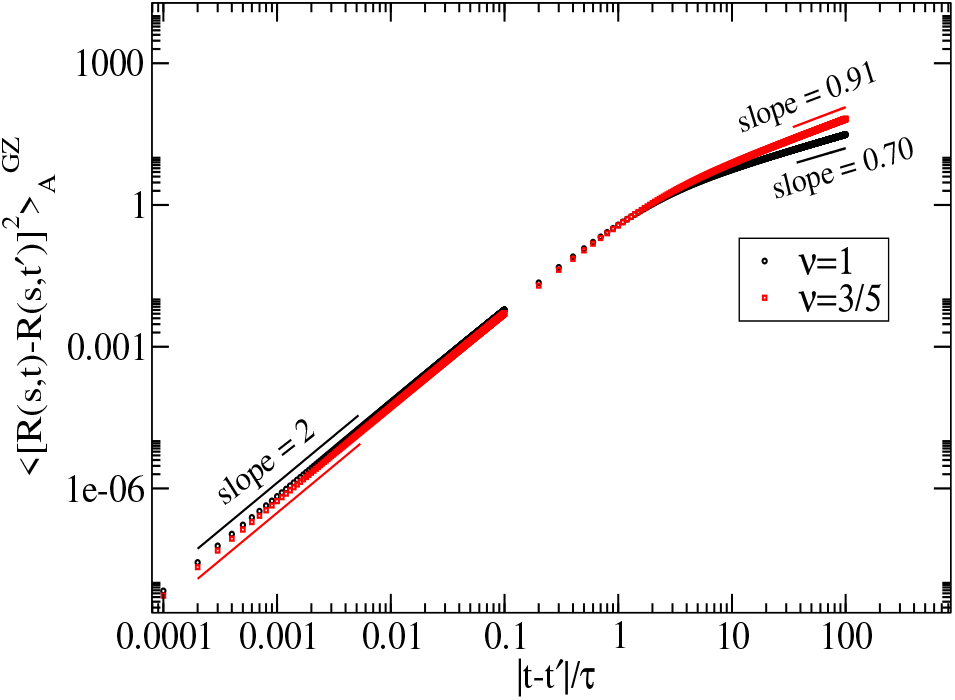
MSD of a segment of Generalised Zimm dynamics with active coupling.

The thermal contribution to MSD from Eq. 23 can be exactly obtained as

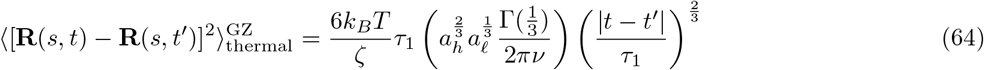

**TABLE II.**
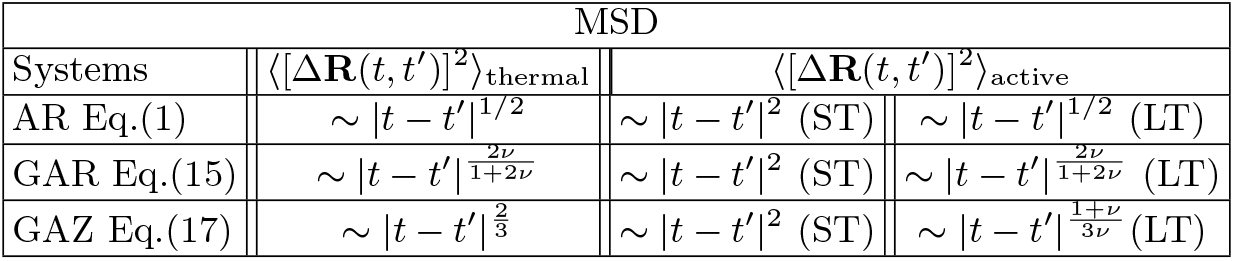
The scaling reltion of MSD for three different dynamics. In the table, ST (|*t* – *t′*| ≪ *τ*) and LT (*τ* ≪ |*t* – *t′*| ≪ *τ_R_*/*τ*_GR_/*τ*_GZ_/) stands for the short-time and longtime limits, respectively.

We study the MSD of a segments for all three models, and we obtain the scaling exponents for short and long-time limits via analytical calculations. We also perform the numerical integration for different *ν* which agree with the analytical results. For short time (|*t* – *t′*| ≪ *τ*), the active contribution to MSD is Newtonian (|*t* – *t′*|^2^) for all models whereas longtime behavior (|*t* – *t′*| ≫ *τ*) coincides with its thermal counterpart except GZ.

The dynamics GAR and GAZ both incorporate segment-segment interactions, whereas GAZ has hydrodynamics interactions in addition to the segment-segment interactions. The model GAZ dynamics can exactly be mapped to GAR for *a_h_* = 1 and *ν* = 1.

## IV. CHARGED MACROMOLECULAR SOLUTION

In this part, we consider a dilute polyelectrolyte solution containing a few homogeneously charged flexible polymer chains, each consisted of *N* number of monomers embedded in a solution of volume *V*. The average concentration of polymer is 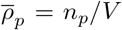 where *n_p_* is the number of polymer chains in the solution. The total charge *a_h_z_p_e* before dispersed in the solution where *e* is the electric charge, and *a_h_* is the degree of ionization. In the solution, we have other species, enzyme-like proteins which are also charged and interact with the charged polymer electrostatically. The total number of enzymes is *n_en_* ≪ *n_p_*. Thus the solution contains three different species, namely the charged polymer and enzymes, and the dissociated counterions, so that the overall system is electrically neutral, i.e,

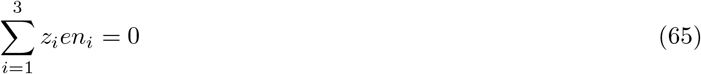

The total number of dissociated counterions is 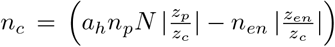 because |*z_c_e*| is the total charge of a single counterion. The dynamics of an enzyme in the solution is driven by the thermal force, screened columbic potentials due to the charged species. In addition, there may be other forces that may come from the chemical reaction, or the shape of the enzymes, which are usually asymmetric, can affect its dynamics. However, the detailed dynamical study of the enzymes is beyond the scope of this work. We assume that the binding and unbinding of an enzyme to polymer is in equilibrium and driven by the electrostatic and thermal force.

Since the density of the enzyme is considered to be very low compared to that of counterions, practically, the enzymes do not affect the counterion’s dynamics. Here we only focus on the dynamics of the macromolecules which get modified by the binding and unbinding of the enzymes and the counterions. In the solution, an enzyme comes near to a polymer and gets attracted due to the electrostatic force and bound to it as illustrated in Fig. (8). Eventually, a bound enzyme gets detached from a polymer due to the thermal fluctuations. As the density of the enzymes is very low, and only an enzyme can bound to a polymer at a given time, thereby any cooperative and competitive effect of the enzymes is negligible.

**FIG. 8.**
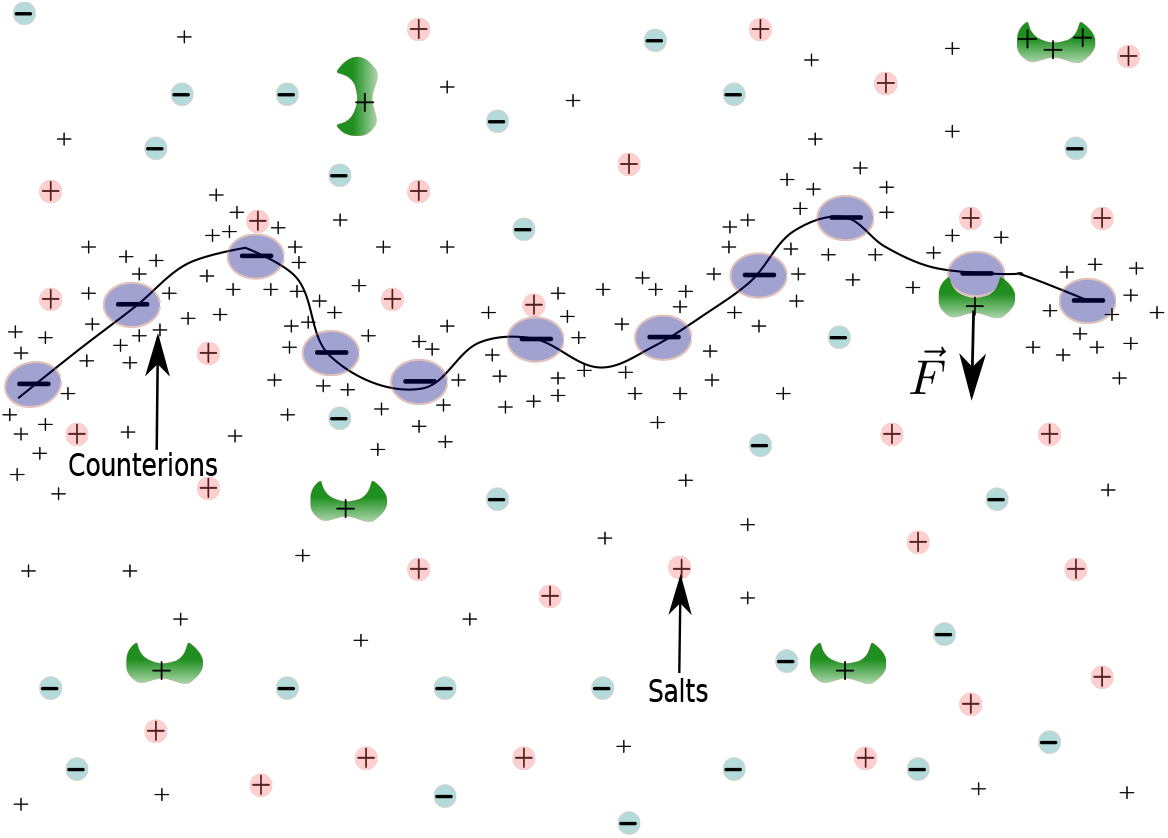
A schematic diagram of a polyelectrolyte solution with the charged enzyme. The macromolecule is negatively charged in the solution. The dissociated counterions are always surrounded by the macromolecule. In addition, salt ions are also present in the solutions.

### A. Cooperative Diffusivity

In an infinitely dilute solution, we consider a polymer chain where s is an continuous variable along the contour of the chain, here write a Langevin equation of a polymer segment with the active coupling where hydrodynamics and segment-segment interactions are taken into account.

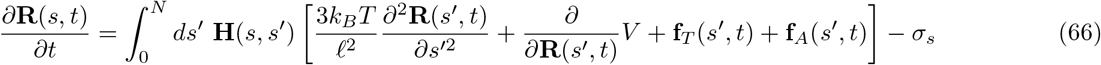

where **R**(*s, t*) is the position vector of a segment of *a_h_*-th chain, and **H**(*s, s′*) corresponds to the hydrodynamic interaction among all the segments. Force experienced by one segment will be carried via **H**(*s, s′*) to other segments. The second term on the right-hand side of Eq. (66) indicate the segment-segment interaction which can be expressed as

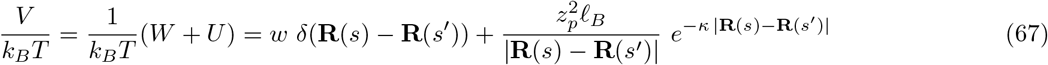

where *w* is the coefficient of the excluded volume interaction and *U* corresponds to the electrostatic interactions. In the above equation 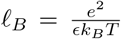 is the Bjerrum length, and *ϵ* is the effective dielectric constant of the solution. The argument *κ* in the exponential of the electrostatic potential is the inverse Debye length which determines the electrostatic screening length due to dissociated counterions and salt ions in the solvent. The last term *σ_s_* is the frictional force due to friction of the solvent to the sth segment.

The hydrodynamic interaction terms **H**(*s, s′*)

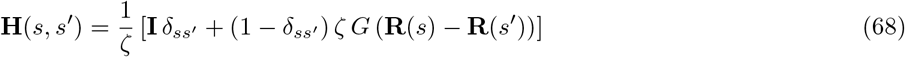

as expressed above is a nonlinear function. Therefore it is very difficult to proceed with the exact form of the term, where ζ in equation 68 is the friction of a segment with the solvent followed from the Stokes-Einstein laws, and *G* (**R**(*s*) – **R**(*s′*)) is the Oseen tensor can be written as

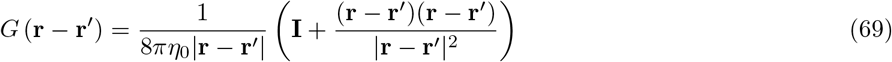

where **I** is the unit tensor. Oseen tensor *G*(**r** – **r**′) acts as a propagator from the point **r** to **r**′. The angular average of 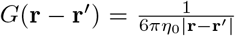. In order to simplify the behavior of **H**(*s, s′*), we follow the preaveraging approximations and uniform expansion expansion due to electrostatic and excluded volume potentials, we obtain

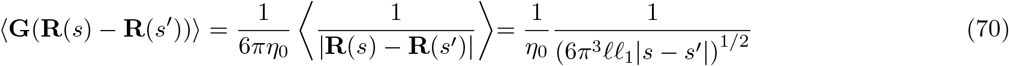

The effect of short-range excluded volume interactions and the segment-segment electrostatic interactions generates a highly nonlinear effect. Assuming the Gaussian chain, the effect of excluded volume interactions and electrostatic interactions can be expressed in terms of effective Kuhn length *ℓ*_1_.

#### 1. Fourier transformation

As shown in the Appendics that the total translational friction on the polymer in the large-scale limit (*q* → 0) obtained as 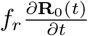. In the absence of the acceleration, this force will be balanced by the thermal noise and the active force applied on the polymer. Assembling all the components in Fourier space in Eq. 66, we write the Langevin equation for the segment as

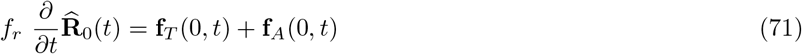

where the effect of solvent, segment-segment interactions are captured by *f_r_* as described in the Appendics. The second and third terms **f**_*T*_(0, *t*) and **f**_*A*_(0, *t*) are the total thermal noise and active force, respectively experienced by the whole polymer. Integrating over all the segments, using Eqs. 2 and 3, we obtain

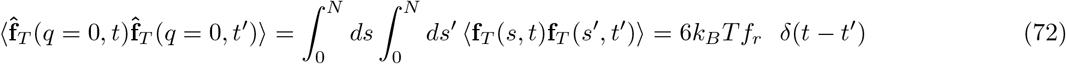

and

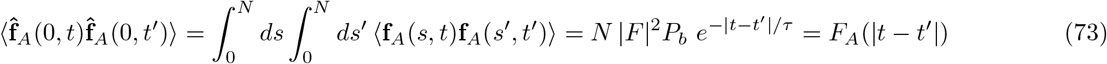

Let us derive the the time evolution of the concentration of the polymer which is derived via Fokker-Planck equation is shown below.

#### 2. Fokker-Planck equation

Let us start with the Stochastic-Liouville equation for the polymer density, which can be written as

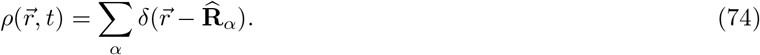

We are interested in the dilute solution, where the interchain entanglement, electrostatic, and hydrodynamics interaction is negligibly small. Incorporating the effect of segment-segment interaction and friction with the solvent, the large scale dynamics simplifies to an effectively single charged polymer with active coupling and its dissociated counterions. Next onwards, we will drop the chain index *α* because of the infinitely dilute solution. The time evolution of the density is obtained as

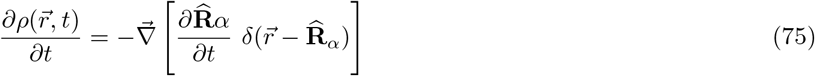

Let 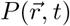 is the probability density of a polymer located at **r**. According to the van-Kampen lemma [13], the probability can be expressed as

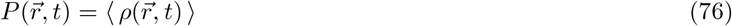

which represents the density in phase space. Substituting Eq. 71 into Eq.75, and considering the ensemble average in Eq. 75, the time evolution of the probability density can be expressed as

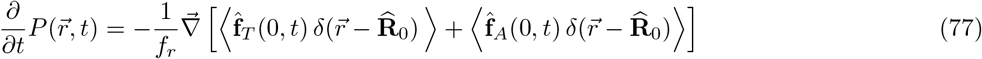

To evaluate the second and third terms on the right hand side of Eq. 77, we employ functional calculus for colored noise [9]. Let us denote functional *F*[*f_T_*] either for thermal noise 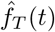 or 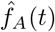, and 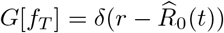 as Eq. 66 contains both the noise terms. The idea is to use the statistical properties [14] of noise *f_T_*(*t*) and *f_A_*(*t*) to calculate the second and third terms on the right hand side in Eq. 77. Assuming the system is in stationary state, and noise is Gaussian, the correlation between two functional can be expressed as

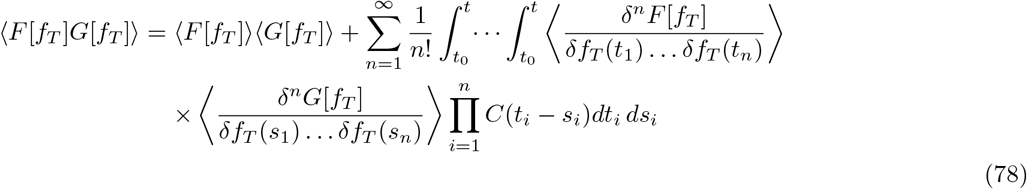

where *C*(*t_i_* – *s_i_*) is the two points temporal correlation function. For this work, in the above equation only *n* = 1 term will contribute, and contribution from all other term *n* ≥ 2 is zero. Using Eq. 78, we obtain the correlation for one component of the thermal noise

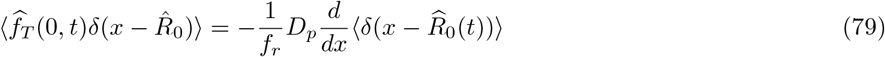

where 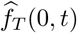 is the Gaussian white noise. For the colored noise, we obtain (see Apendix for the detailed calculation)

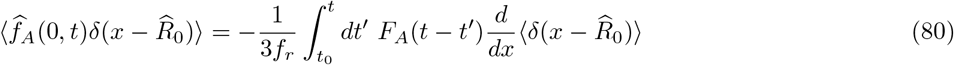

where *F_A_*(|*t* – *t′*|) is the autocorrelation of the active force given in Eq. (3). Hence, assuming the stationary state Gaussian noise, we obtain a Fokker-Planck equation which is expressed as

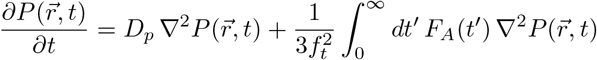

where 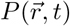 is the probability density of the center of mass of the polymer, and 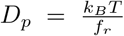, and *F_A_*(|*t′*|) = *NP_b_*|*F*|^2^*e*^−|*t*′|/*τ*^. In the dilute solution, due to the hydrodynamics interaction, the translational diffusion coefficient in the above equation follows Zimm dynamics and inversely proportional to the radius of gyration of the polymer [10–12].

In the absence of the excluded volume and electrostatic interaction, we have 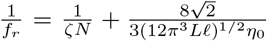. Using the screened excluded volume and the electrostatic interaction, Kuhn length *ℓ* gets modified to *ℓ*_1_, which lead the above equation to

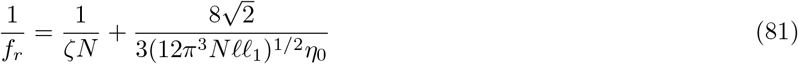

In the free draining limit, the diffusivity 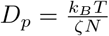. When the hydrodynamic interaction is dominant

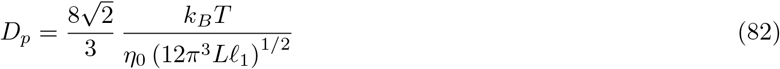

where *η* is the diffusivity of the solution and *R_g_* is the radius of gyration of the polymer. The active coupling of the enzymes on the polymer give rise to the effective diffusivity which is obtained as where

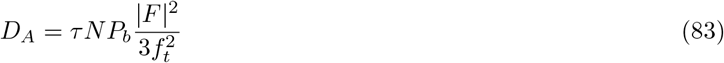

where

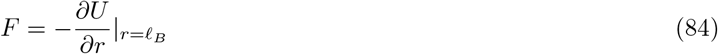

where the potential is

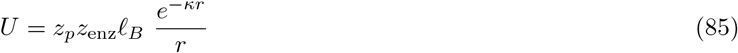

The magnitude of the force can be obtained as

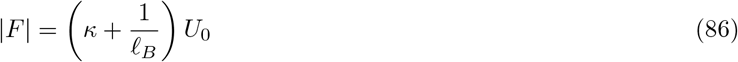

where *U*_0_ is defined at a Bjerrum length as

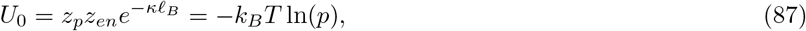

since *U*_0_ = –*k_B_T*ln(*p*). Hence, |*F*|^2^ can be expressed as

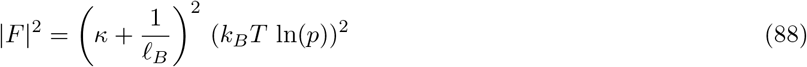

In low salt limit *κ* → 0, 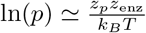, hence the magnitude of the force |*F*| can be obtained as

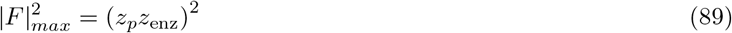

In high salt limit *κ* → ∞, ln(*p*) → 0, hence the magnitude of the force |*F*| = 0. The binding and unbinding timescale can be expressed in terms of *p* which yields

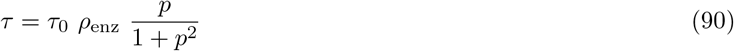

Substituting Eqs. 90, 88 81, and in Eq. 83, we obtain the expression for the diffusivity due to active coupling as

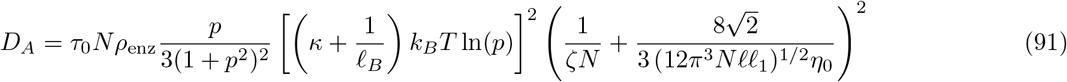

This shows how active part of the diffusivity *D_A_* explicitly depends on various things as *κ, N* and effective Kuhn length *ℓ*_1_. The expression *ℓ*_1_ [33, 38] which captures the effect of the electrostatic interactions between the segments and the effect of the excluded volume as

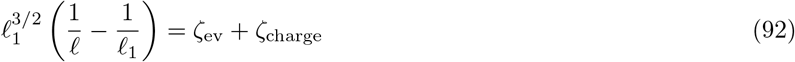

where

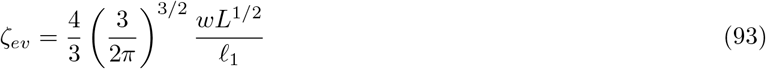

and

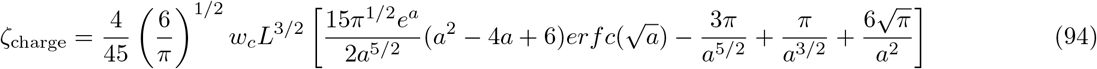

where 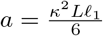. This show that the dependencies of effective Kuhn length *ℓ*_1_ on *κ* is very nontrivial.

### B. Counterions coupling

In this subsection we consider the coupled dynamics of the counterions and derive the cooperative diffusivity of the polymer in infinitely dilute solution. Since a charged polymer in a solution is always surrounded by its dissociated counterions clouds, the dynamics of counterions and polymer are always coupled and get significantly modified by the coupling of the counterions. Though the overall solution is electrically neutral, the presence of the counterions gives rise to density fluctuations and induces local electric field 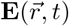 which affect the dynamics of the polymer. In the absence of active coupling, when the charged polymeric solution is infinitely dilute, the scenario becomes very simple and can be described by well known Nernst-Hartley theory [17]. This leads Eq. 81 to take an equation of continuity as

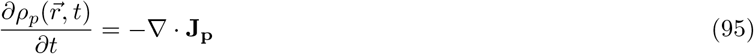

where 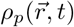 is the local polymer concentration, and current

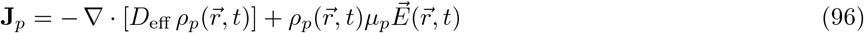

and the second term in Eq. 96 is the convection term from the density fluctuation of the local charged species.

The dynamics of the concentration of counterions is strongly coupled to the dynamics of the concentration of the polymer. The equation of motion of the counterions can be written as

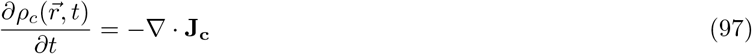

where *ρ_c_*(*r, t*) is the local counterionss concentration, and current

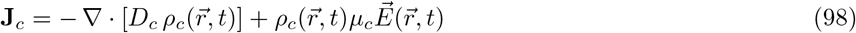

These three species are dynamically very different, and since time dependent concentration fluctuations of the species can be measured in the dynamical light scattering experiment, we express concentration fluctuations up to linear order as

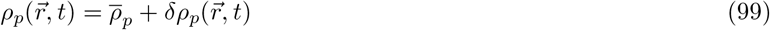

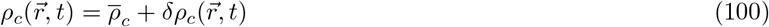

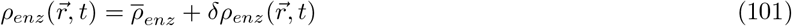

where 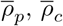, and 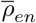 are the average concentration of polymer, counterions and charged enzymes, respectively.

The divergence of the electric field can be written as the Poisson equation

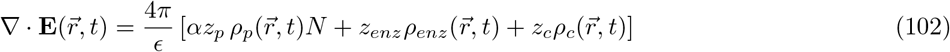

The density of the enzyme is considered to be very low *ρ_enz_* ≪ *ρ_c_*. The effect of the binding of the enzyme on the polymer because of the electrostatic interactions which has already been considered in the time evolution equation for the polymer concentration.

In order to study the coupled dynamics as given by Eqs. 95, 96, 97, 96 in the leading order approximations, here we define the Fourier transformation of 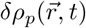 as

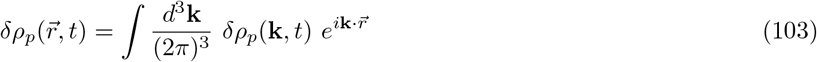

where **k** is the scattering wave vector. Here we obtain the time evolution equation of the small fluctuations in the concentration field of the polymer and counterions in which we only interested in fluctuations up to linear order[]. Using the electroneutrality, the coupled equations becomes in Fourier space as

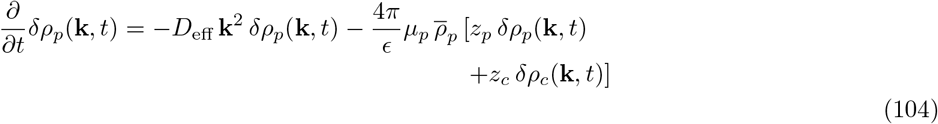

similarly for counterions

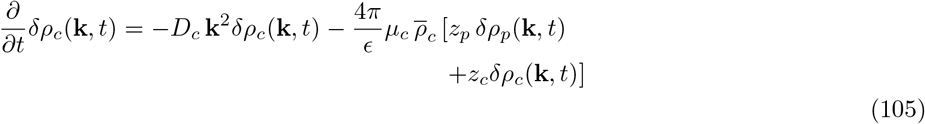

where *ϵ* is the dielectric constant of the solution. 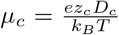, and similarly 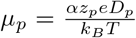. The relaxation time scale of the counterions are much faster compared to macromolecules. Therefore the fluctuation of polymer concentration experience the steady state behavior of the counterions, i.e., 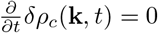.

### C. Results and Discussions

Considering the expression of *δρ_c_* in steady state, we get the expression for *δρ_c_* from Eqs. 105 and substitute it in 104, we obtain where

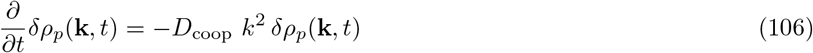

where

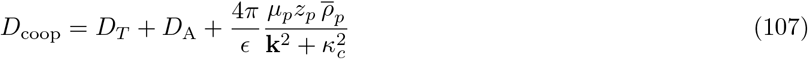

where

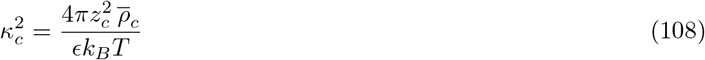

Substituting *κ_c_* from Eq. 108 and *μ_p_* from Eq. (135), and also *D_T_* = *k_B_T/f_r_* into Eq. (110), we obtain

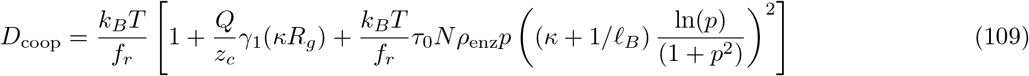

we also use *Q* = *a_h_z_p_eN*, and 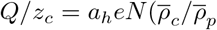, which yields

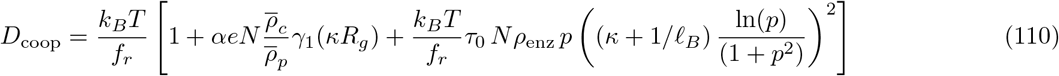

Thus from Eq. 106, we can obtain time correlation function which is obtained as

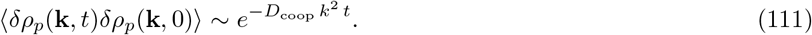

In Eq. (111), the density fluctuation can be measured in the dynamic light-scattering where *k* is the scattering wave vector.

## V. SUMMARY AND CONCLUSIONS

In this work, we have introduced an active coupling due to the binding and unbinding of the enzymes, as the enzymes carry charge and bind to a segment electrostatically. We first start with the dynamics of a segment and its segment-segment correlation for a simple Rouse model with active coupling. We further study two other dynamical models of polymers, namely generalized Rouse, and generalized Zimm model which incorporate segment-segment interactions without hydrodynamics interactions and segment-segment interactions with hydrodynamics, respectively. In the three models, we systematically study the segment-segment correlation, MSD of each models and compare the results from the thermal and active contribution of each model, and we also compare the same quantity (e.g., MSD) among the models.

Finally, we derive a theory for the dynamics of the concentration fluctuations of the macromolecules with hydrodynamics and segment-segment interactions wherein the dynamics of macromolecules are coupled with the counterions and also driven by active coupling in an infinitely dilute solution. We use the translational friction coefficient and mobility of macromolecules and derive the continuity equation for the polymer concentration with active coupling in a dilute solution. We also couple the dynamics of the counterions with polymer concentrations and considering density fluctuations up to linear order, we derive the cooperative diffusivity of the macromolecular solutions in infinite dilute limits. Our obtained closed-form expression for the diffusivity of the macromolecules demonstrates that how the activity-induced diffusivity depends on the degree of polymerization, degree of ionization, temperature, charge density of the macromolecules, counterions, and enzymes along with the other transport properties of the counterions and polymers.

On the other hand, the expression for the diffusivity does not consider the detailed dynamics of the enzymes, namely shape, conformational change of the enzyme, and also not consider the conformational change of macromolecules during binding. These details of the enzyme’s dynamics may be crucial in a real-life biological system.

For the equal-time segment-segment correlations and MSD, though we consider the electrostatic binding as the source of active coupling, we express our results in terms of |*F*| which is general wherein any type of local force may cause the colored noise, and can be treated in this approach. Importantly, we also express the segment-segment correlation and MSD in terms of size exponent *ν*, therefore, for any feasible size exponent, the calculation and the obtained scaling relation along with its coefficient are valid. Since the calculation involves Fourier mode *q* corresponding to the polymer which is always one dimensional, the obtained scaling form remains the same for other dimensions, only the size exponent *ν* would be different depending on the dimensions.

Here we open up an active coupling due to electrostatic attractive force which is ubiquitous in biological systems and its application in biomedical and synthetic bio-polymeric systems. To understand a real-life biological system, namely cell, the environment is very crowded and the solution is also salty. In the absence of active coupling, in a simple polyelectrolyte solution, added salt concentration takes a very important role in the dynamics of this charged macromolecule. Because with the increase of the concentration of salt in the solution, the electrostatic screening will be increased, and consequently, the dynamics become significantly different. In this view, with the active coupling, the effect of added salt on the mode of relaxation would be an important future work both in the dilute and high concentration limits of the polymers.

## ACKNOWLEDGMENT

Acknowledgement is made to the National Institutes of Health (Grant No. 5R01HG002776-16), the National Science Foundation (DMR-2015935), and the AFOSR Grant FA9550-20-1-0142 for financial support.

## APPENDIX

## CALCULATION OF THE ACTIVE CONTRIBUTION TO MSD: ACTIVE ROUSE MODEL

Here we present the calculation of the active contribution to the MSD for active Rouse model. The calculation for GR, and GZ dynamics would also be the similar.

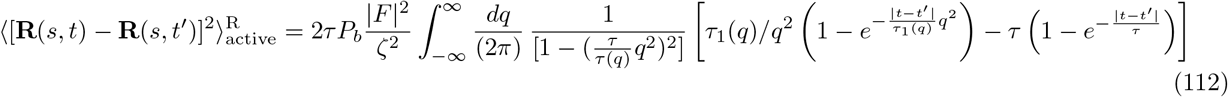

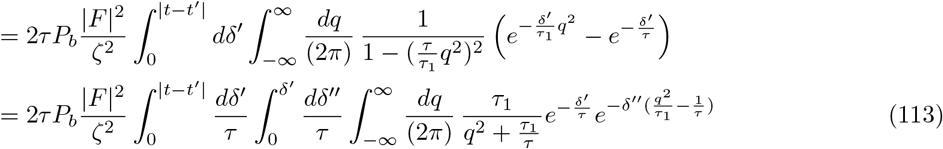

In the above equation, we swap the integration of *δ′* and *δ″* without chainging the generality we have

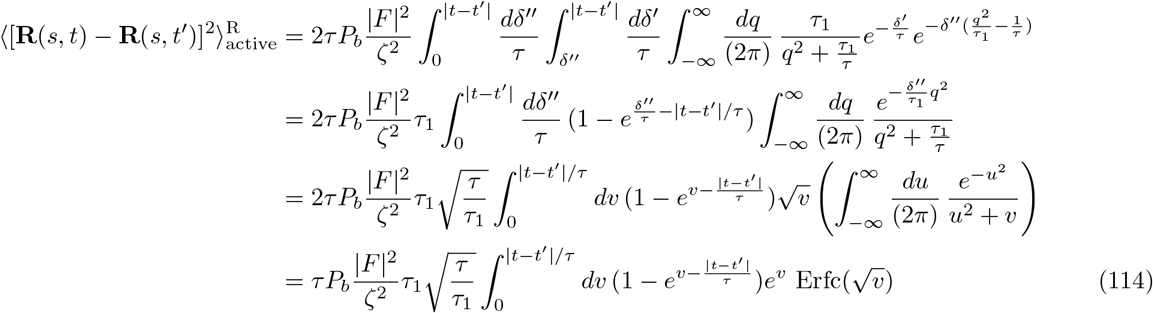

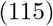

where we used

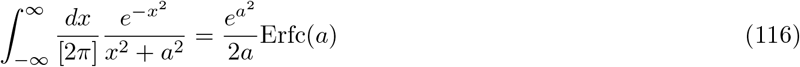

The active contribution calculation for other dynamics, generalised active Rouse and Generalised-active Zimm will also be the same. One useful integration results is given which is useful for the calclation of MSD of other dynamics

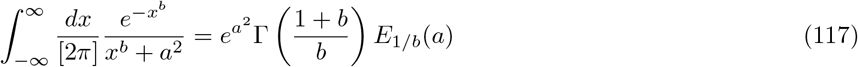

where

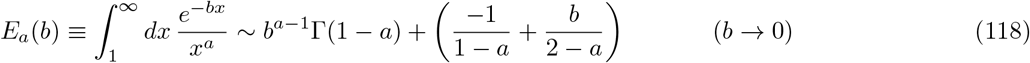

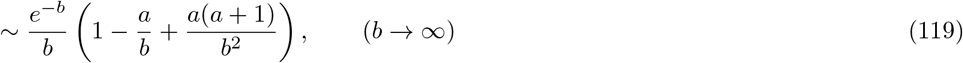

## OVERVIEW OF THE CALCULATION OF FRICTION COEFFICIENT OF CM OF A POLYMER AND ELECTROPHORETIC MOBILITY

### A. Solvent

In very low Reynolds number limits in the absence of polymer molecules, the Navier-Stokes equation for the solvent can be described as

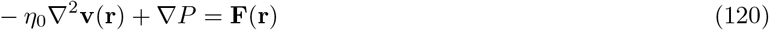

where *η*_0_ is the shear viscosity, *P* is the pressure in the solvent, and **F**(**r**) is the force field in the solvent. Imposing incompressibility condition on the solvent, i.e., ∇ · **v**(**r**, *t*) = 0, we eliminate the pressure term in Eq. 120, and it leads to

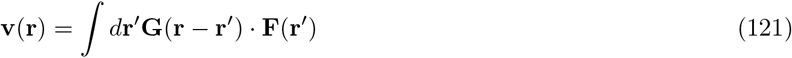

where *G*(**r** – **r**′) is the Oseen tensor.

### Polymer and Solvent

In the presence of the polymer in the solution, the dynamics of the solvent and polymer get coupled to each other. In this scenario, the *i*th segment of a polymer exerts force *σ_i_* on the fluid and thereby it experiences force –*σ_i_*.

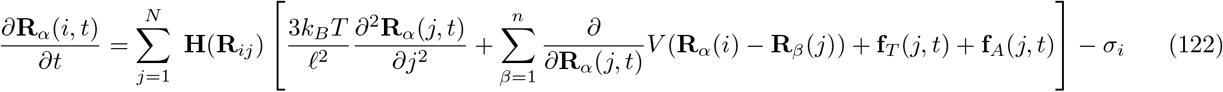

The equation for the solvent can be expressed as

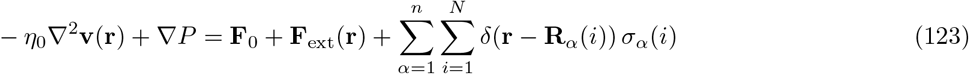

where force **F**_0_ causes the velocity **v**_0_(*r*) in the absence of the polymer. The velocity of the solvent where *α* is the index of the chain. Following Eq. 121, the velocity field for the solvent in the presence of the polymer can be expressed as

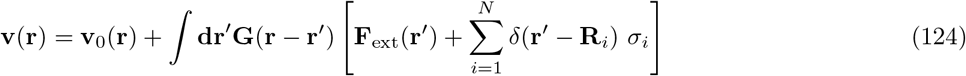

Using the no-slip boundary condition at **R**_*i*_, the motion of the solvent and the motion of the segment can be equated as

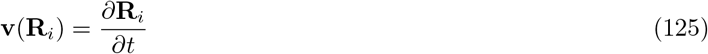

Using Eq.125 in Eq. 124, we express the velocity of solvent in terms of velocity of segments as

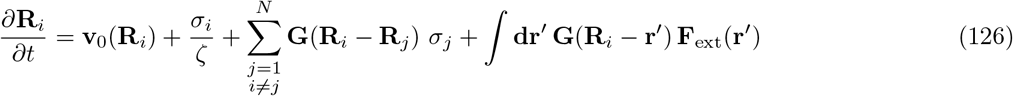

Averaging over chain configurations, we consider **v**_0_(**R**_*i*_) = 0 and **F**_ext_ = 0, we obtain

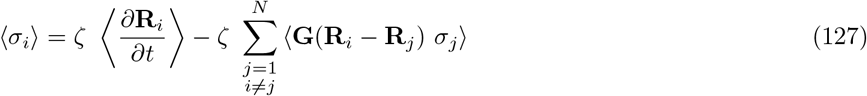

At this point, in order to solve the above equation a couple of assumptions are employed. Firstly, the drift velocity of each segment is the same and that is equal to the CM of the macromolecules. In Eq. 127, the second term represents that magnitude of the force *ζ σ_j_* experienced by *j*th segment and propagation of the force from *j*th to *i*th segment. Secondly, since the exact solutions are very difficult, and it is wisely employed that assumptions [?] that on conformational average the force experienced by each segment *σ_j_* is same in the solvent which leads to decouple the force *σ_j_* and Oseen tensor *G*(**R**_*i*_ – **R**_*j*_). This simplifies the equation as

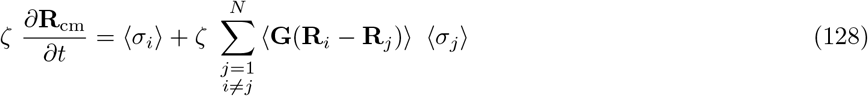

Here another assumptions, namely the segment-segment electrostatic and excluded volume interactions lead uniform expansion of the chain. Substituting these in the above equation and solving the integral equation [39]. The frictional froce experienced by the whole chain is 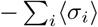. In an infinitely dilute solutions, balancing the force the motion of CM can be written as

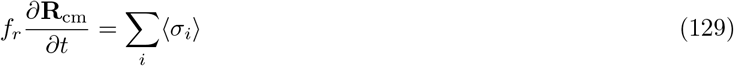

Using Eq. 129 and 70, and substituting these in Eq. 128, the expression of *f_r_* is derived as

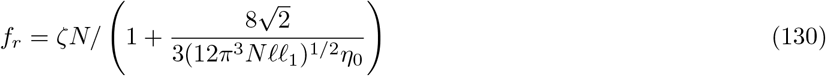

Since the friction coefficient *f_r_* of the CM the whole polymer chain the translational friction coefficient [37, 40] can be obtained as

In deriving the translational friction coefficient, the segment-segment interactions and the effect of solvent is included in a nontrivial way.

### Calculation of electrophoretic mobility

A homogeneously charged polymer in a solution under the applied constant electric field *E* moves with a drift velocity following the relation

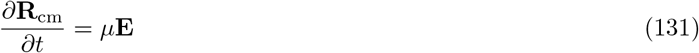

where *μ* is the electrophoretic mobility of the polymer ia an infinitely dilute solution. In the absence of acceleration of the polymer, the total force on the polymer due to the electric field is exactly balanced by the frictional force which suggest

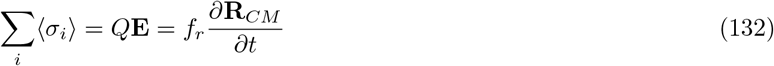

where *Q* = *Nz_p_e* is the total charge of the polymer and *z_p_e* is the charge of a single segment. As we know that a charged polymer in a solution is always surrounded by its dissociated counterions. Around each segment of the polymer, there is a opposite charge counerion cloud. This will induce an electric field which affect the segments and each segment wll affect the other segments in the presence of the hydrodynamics interactions. The force field **F**(**r**) from the potential field from the counterions will appear as a force as

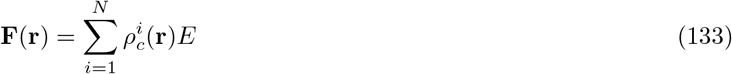

where 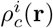 is the density of the counterions due to *i*th segment. Note that at position **r**, the counterions of many segments may contribute to give rise the force which is captured by the 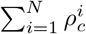. Because of the electroneutrality, the counterions density around an isolated segment can be defined as

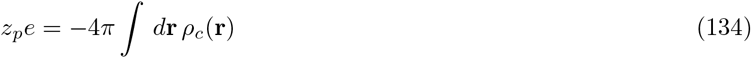

Incorporating the effect of counterions in the coupled equation of the solvent and polymer, and assuming the same approximation, namely uniform expansion and averaging approximations, the mobility can be derived as

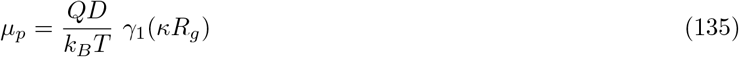

where *κ* and *R_g_* are the inverse Debye length and radius of gyration of the polymer. The factor *γ*_1_ comes due to the effect of the counterions and the approximate expression is also complicated. One scaling limit s very simple to note that when *κ* → 0, *γ*_1_ = 1.

### Calculation of Correlation for Functionals

In this Appendix, by following Hänggi’s prescription for the Fokker-Planck equation with colored noise, we derive the concentration equation for the polymers in a infinitely dilute solution given by Eq. (66). Let us consider a functional *G*[*η*(*t*)+*x*(*t*)] where *x*(*t*) is a test function of *η*(*t*). The Taylor series expansion of *G*[*η*(*t*)+*x*(*t*)] in *η*(*t*) can be expressed as

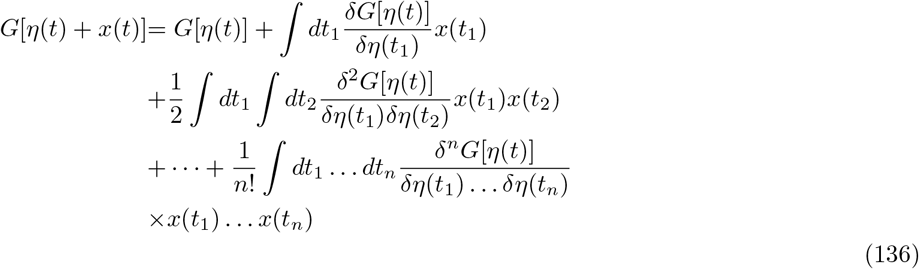

When the noises are assumed to be Gaussian, the statistical correlation between any two arbitrary functional can be expressed as

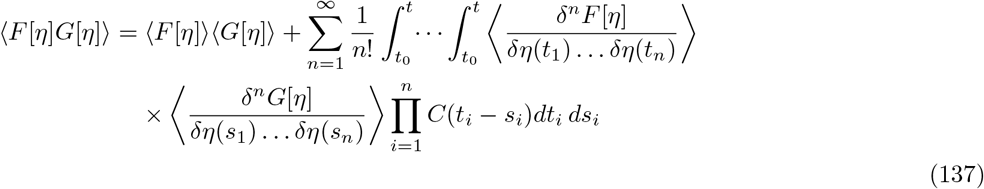

where *C*(*t_i_* – *s_i_*) = 〈*x*(*t_i_*)*x*(*s_i_*)〉 is the second order cumulant.

### Calculation for white Noise

Considering, the *F*[*η*] = *x*(*t*) in Eq. 137, we obtain

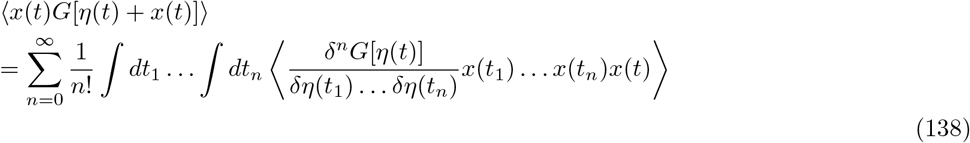

which can be expressed as

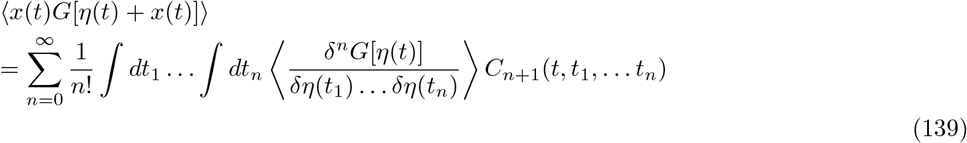

It is better to deal the variable as scalar, therefore we consider one component of force 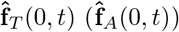 i,e., 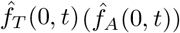 and carry out the calculation. Thereafter we generalize for three dimensions without changing the generality. Let us first start with the thermal noise correlation 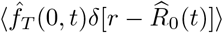, and active noise correlation 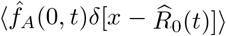, in Eq. 77. We obtain

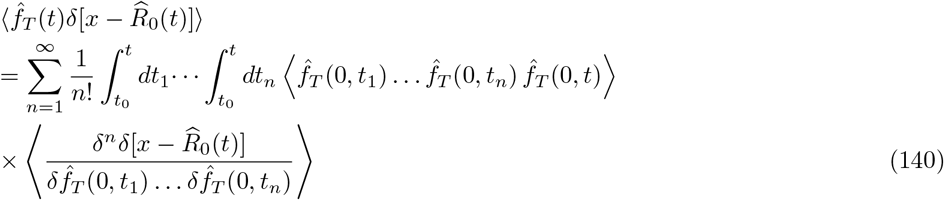

Using Eq.72 the statistical properties 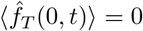, and 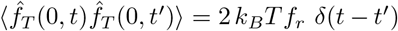 for one component. In the above equation, only the term *n* = 1 contributes and terms *n* ≥ 2, vanishes. Thus we now get,

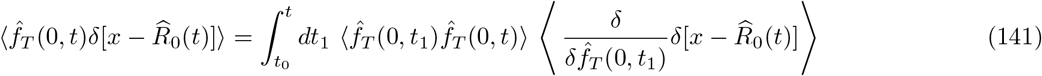

Here, we can write,

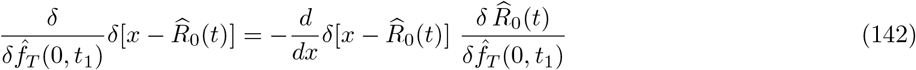

Using the Langevin equation described in Eq. 66, as the thermal noise 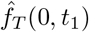 is linearly proportional to **R**, we obtain

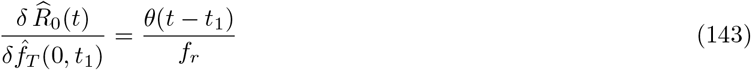

Substituting the above equations in Eq. 142, we obtain

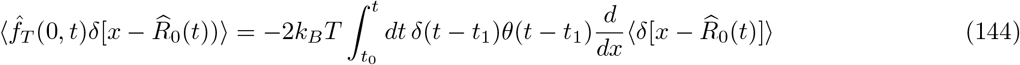

where 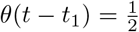 if *t* = *t*_1_. [29] We have now obtained

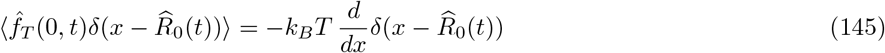

### Calculation for Active Force

Following the above shown prescription for thermal noise, we can get

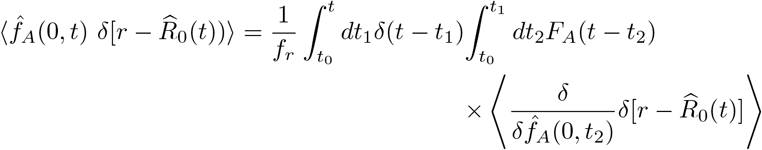

where functional differentiation of the delta function can be obtained as

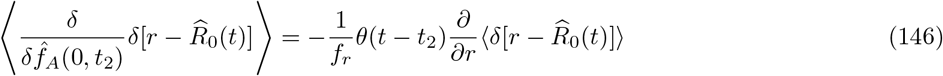

Substuting Eq. 146, in Eq. 146, we obtain

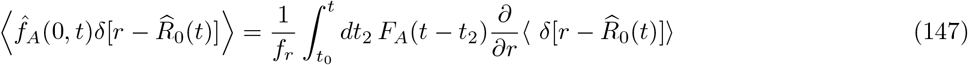

where *t*_0_ is set to be zero, and in steady state time *t* → ∞ is considered.

## Notes

### Competing Interest Statement

The authors have declared no competing interest.

